# Evolution in interacting species alters predator life history traits, behavior and morphology in experimental microbial communities

**DOI:** 10.1101/748582

**Authors:** Johannes Cairns, Felix Moerman, Emanuel A. Fronhofer, Florian Altermatt, Teppo Hiltunen

**Affiliations:** Wellcome Sanger Institute, Cambridge, CB10 1SA, UK; Organismal and Evolutionary Biology Research Programme, Department of Computer Science, 00014 University of Helsinki, Finland; Department of Microbiology, P.O. Box 56, 00014 University of Helsinki, Finland; Department of Aquatic Ecology, Eawag, Swiss Federal Institute of Aquatic Science and Technology, Überlandstrasse 133, 8600 Dübendorf, Switzerland; Department of Evolutionary Biology and Environmental Studies, University of Zurich, Winterthurerstrasse190, 8057 Zürich, Switzerland; ISEM, University of Montpellier, CNRS, EPHE, IRD, Montpellier, France; Department of Biology, 20014 University of Turku, Finland

**Keywords:** predator-prey interactions, trait evolution, ciliate physiology, microbial model systems, experimental evolution

## Abstract

Predator-prey interactions are key for the dynamics of many ecosystems. An increasing body of evidence suggests that rapid evolution and co-evolution can alter these interactions, with important ecological implications, by acting on traits determining fitness, including reproduction, anti-predatory defense and foraging efficiency. However, most studies to date have focused only on evolution in the prey species, and the predator traits in (co-)evolving systems remain poorly understood. Here we investigated changes in predator traits after ~600 generations in a predator-prey (ciliate-bacteria) evolutionary experiment. Predators independently evolved on seven different prey species, allowing generalization of the predator’s evolutionary response. We used highly resolved automated image analysis to quantify changes in predator life history, morphology and behavior. Consistent with previous studies, we found that prey evolution impaired growth of the predator. In contrast, predator evolution did not cause a clear increase in fitness when feeding on ancestral prey. However, predator evolution affected morphology and behavior, increasing size, speed and directionality of movement, which have all been linked to higher prey search efficiency. These results show that in (co-)evolving systems, predator adaptation can occur in traits relevant to offense level without translating into an increased ability of the predator to grow on the ancestral prey type.

## Introduction

Predator-prey interactions are ubiquitous and determine the dynamics of many ecosystems. Predation has been widely studied at an ecological level [1–3], and recent research also shows that this interaction can be strongly altered by rapid evolution of anti-predatory defense in the prey [4] as well as by counter-adaptations in the predator [5–7], even though selection may be asymmetric resulting in slower evolutionary change for the predator [8]. Moreover, owing to population growth-defense tradeoffs, rapid evolution of the prey and adaptation to predation can result in frequency-dependent selection of defended and undefended prey types as a function of predator population size [9–11], an example of eco-evolutionary feedback dynamics. Common to this spectrum of evolutionary, co-evolutionary and eco-evolutionary dynamics is that these dynamics are all driven by natural selection acting on fitness-relevant traits.

Predation can be described by three main phases, namely prey search, capture and ingestion [12]. These three phases are shaped by key traits in predator-prey systems, including those influencing offence and defense level, and all these traits can be subject to evolutionary change [13]. Offense level is determined by sensory faculties and speed enabling location and capture of prey, and defense level by the capacity for predator avoidance and escape prior to ingestion as well as physicochemical obstruction of ingestion and digestion. Adaptations in defense and offense, in turn, combined with associated tradeoffs, modulate the reproduction (i.e. life history traits) of both parties. Examples abound of the study of the different phases of predation, and adaptation in both predator and prey life history traits. For example, the timing and population dynamics of many insectivorous bird species are tightly coupled to the dynamics of their prey insect species [14]. Olive baboon sleeping site choice and behavior (sharing sleeping sites between multiple baboon groups) in Kenya were recently linked to decreased contact and capture rate by leopards [15]. Co-evolution has been hypothesized to occur between Northern Pacific rattlesnakes and California ground squirrels whereby venom resistance in squirrels is matched by increased venom effectiveness in rattlesnakes based on field data supportive of local adaptation of the traits [16]. All of these empirical examples are, however, limited to a mostly comparative and behavioral ecology study approach, and cannot be used to experimentally investigate (co-)evolution in predator-prey systems due to the long generation times of the species.

Microbial systems offer a unique opportunity to study predator-prey dynamics, as they include efficient predators and allow for high replication as well as experimental approaches capturing both ecological and evolutionary dynamics. Microbial predator-prey systems show many key characteristics found also in other predator-prey systems, such as offense by speed [17] and defense by avoidance of detection [18], escape [19], or physicochemical obstruction of ingestion or digestion (for an overview, see [12]). Defense level has also been demonstrated to evolve in controlled setups [20, 21]. However, to our knowledge, there exist little to no empirical studies examining offense mechanisms subject to rapid evolution in microbial predator-prey systems.

Here we employed an experimental evolution approach to test the influence of ~600 generations of predator-prey interaction on predator traits, using a microbial (ciliate-bacteria) model system. Since predator-prey dynamics are characterized by the intrinsically linked dynamics of both interaction partners, we inspected the influence of both prey and predator evolution on predator traits. To find general patterns in predator traits independently of any specific prey species, as most predators have multiple prey species, we used seven different prey species that were all separately evolved with the predator. Predator traits, analyzed using high-resolution video recording, aligned with theory on physiological factors increasing prey search efficiency for morphological and behavioral trait evolution [17, 22], although the patterns were often observed only for a subset of species. In contrast, life history traits showed both expected and unexpected patterns, such that prey evolution impaired predator growth as predicted while predator evolution failed to influence growth on the ancestral prey type [7, 23].

## Material and methods

We studied the evolutionary dynamics of one focal predator species (the ciliate *Tetrahymena thermophila*) and seven of its bacterial prey species in all seven combinations of predator-prey species communities, as well as dynamics in prey-species populations only. Predator-prey dynamics and (co-)evolution are relatively specific but less constrained than host-pathogen dynamics, which often only contain a 1:1 species match while predators can frequently feed on multiple different prey species [24]. We ran predator-prey evolutionary experiments over about 600 predator generations, and assessed evolutionary effects on life history, morphology and behavior using common garden experiments.

### Strains and culture conditions

The seven prey species used in this study are listed in Table 1. In addition to four taxa previously used as models in predator-prey studies, three strains were chosen based on representing genera associated with ciliate predators in natural habitats or potentially exhibiting different anti-predatory defense mechanisms (Table 1). Since all the strains represent unique genera, they are referred to by their genus name in the text. As a generalist predator, capable of consuming all the prey species, we used a single strain of the asexually reproducing ciliate *Tetrahymena thermophila* 1630/1U (CCAP) [25].

**Table 1.** Bacterial strains used in this study.

| Strain* | Rationale for species selection |
| --- | --- |
| *Escherichia coli* ATCC 11303 | model prey [26] |
| *Janthinobacterium lividum* HAMBI 1919 | pre-/post-ingestion defense: toxin release [12] |
| *Sphingomonas capsulata* HAMBI 103 | model prey [27] |
| *Brevundimonas diminuta* HAMBI 18 | realistic habitat [28] |
| *Pseudomonas fluorescens* SBW25 [29] | model prey [30] |
| *Comamonas testosteroni* HAMBI 403 | pre-ingestion defense: oversize [12] |
| *Serratia marcescens* ATCC 13880 | model prey [27] |

\*ATCC = American Type Culture Collection; HAMBI = HAMBI mBRC = Microbial Domain Biological Resource Centre HAMBI, University of Helsinki, Finland

Prior to the experiments, all bacterial stocks were kept at −80 °C and ciliate stocks were cultured axenically in proteose peptone yeast extract (PPY) medium containing 20 g of proteose peptone and 2.5 g of yeast extract in 1 L of deionized water. During the evolutionary experiment, cultures were kept at 28 °C (± 0.1 °C) with shaking at 50 r.p.m.

### Predator-prey evolutionary experiment

The evolutionary experiment was started using a small aliquot (20 *μ*L) of a 48-h bacterial culture started from a single colony and 10,000 ciliate cells (approx. 1,700 cells mL^−1^) from an axenic culture. Each bacterial strain was cultured alone and together with the ciliate predator (three replicates each, with the exception of six replicates for *Comamonas*) in batch cultures of 20 mL glass vials containing 6 mL of 5 % KB medium, with 1 % weekly transfer to fresh medium.

Every four transfers (28 days), bacterial and predator densities were estimated using optical density (1 mL sample at 600 nm wavelength) as a proxy for bacterial biomass and direct ciliate counts (5 × 0.5 *μ*L droplets using light microscopy) as used in this context and described previously [30–32], and samples were freeze-stored with glycerol at −20 °C for later analysis. Since predators do not survive freeze-storage in these conditions, at time points 52 and 89 weeks, predator cultures were made axenic by transferring 400 *μ*L into 100 mL of PPY medium containing an antibiotic cocktail (42, 50, 50 and 33 *μ*g mL^−1^ of kanamycin, rifampicin, streptomycin and tetracycline, respectively) and stored in liquid nitrogen. Axenicity was controlled for by plating on agar plates containing 50 % PPY medium where all experimental bacterial strains grow. The liquid nitrogen storage protocol was modified from a previously used protocol [33] and included starving a dense ciliate culture in 10 mM Tris-Hcl solution (pH 7) for 2-3 days, centrifugation (1700 g, 8 min, 4 °C), resuspension of the pellet in 1 mL of leftover supernatant, and the addition of 4 mL of sterile 10 % DMSO. The resultant solution was transferred to cryotubes in 0.3 mL lots, and frozen in a −20 °C freezer at a rate of −1 °C/minute using a Mr. Frosty™ Freezing Container (Thermo Scientific) for cell preservation before transferring to liquid nitrogen.

### Sample collection and preparation

We isolated the populations for the current experiment at time point 89 weeks (approx. 20 months). With the minimal assumption that populations multiply by 100-fold (dilution rate) until reaching the stationary phase, each weekly transfer interval represents 6.64 generations for both prey and predator [34], constituting a total minimum of ~600 generations. Community dynamics are shown in Supplementary Figures S1 and S2 and show clear differences in population size between different prey species.

Bacteria were restored from freeze-storage by transferring 20 *μ*L into 5 mL of 5 % KB medium and culturing for 72 h. Predators were restored from liquid nitrogen by thawing cryotubes in a 42 °C water bath for 15 s, followed by the addition of 1 mL of 42 °C PPY medium. The cryotube contents were then transferred a petri dish containing PPY medium at room temperature. Upon reaching high density (approx. 48 h), predators were transferred to 100 mL of PPY medium and cultured to high density (approx. seven days). To ensure that the antibiotic treatment or the liquid nitrogen storage and revival procedures do not contribute to potential differences between the ancestral predator and evolved predator lines, the axenic ancestral predator was subjected to identical procedures and was revived at the same time as the evolved lines. These culturing steps representing over 10 generations should remove the influence of non-genetic changes in predator traits caused by phenotypic plasticity [35].

### Physiological measurements

To test bacterial and ciliate performance and traits, we used a combination of automated video analysis, optical density measurements and flow cytometry. To separate evolutionary responses on the predator and prey level, we tested performance of both evolved and ancestral bacteria with evolved and ancestral ciliates for all evolved lines reciprocally. To do so, we prepared 12 50 mL falcon® tubes by adding 20 mL of 5 % KB medium. Three of these were inoculated with ancestral bacteria and ancestral ciliates, three with ancestral bacteria and evolved ciliates, three with evolved bacteria and ancestral ciliates and the remaining three with evolved bacteria and evolved ciliates. We placed the falcon® tubes in a 28 °C incubator, rotating on a shaker at 120 r.p.m. After inoculation, the samples were left to grow for a period of 12 days, to allow populations to grow to equilibrium density. Over the course of these 12 days, we took a total of 10 samples from each culture for analyzing population density dynamics of bacteria and ciliates, and morphological and behavioral metrics for the ciliates.

### Bacterial density measurements

Bacterial density was determined through measurement of both optical density and through flow cytometry. Flow cytometric analyses, were based on established protocols [36, 37] which facilitate distinction between living bacterial cells and background signals (e.g. dead cells or abiotic matter). For flow cytometry, we sampled 50 *μ*L of all cultures, diluted 1:1000 using filtered Evian water and transferred 180 *μ*L of the diluted samples to a 96-well-plate. We then added 20 *μ*L of SybrGreen to strain the cells and measured bacterial cell counts using a BD Accuri™ C6 flow cytometer. As the inner diameter of the needle from the flow cytometer was 20 *μ*m, and hence smaller than typical ciliate cell sizes, it is highly unlikely that ciliate cells were accidentally measured during flow cytometry. Also, given that bacterial densities were typically between one to five orders of magnitude larger than ciliate densities, even an occasional measurement of ciliate cells would have a negligible effect on bacterial density estimates. The full protocol can be found in Supporting Information. For optical density measurement, we sampled 50 *μ*L of all cultures, diluted 1:10 using filtered Evian water, and measured absorbance at 600 nm using a SpectroMax 190 plate reader.

### Ciliate density and trait measurements

For measuring ciliate density, we used a previously established method of video analysis [38] using the BEMOVI R-package [39]. We here followed a previously established method [40] where we took a 20 s video (25 fps, 500 frames) of a standardized volume using a Leica M165FC stereomicroscope with circular lighting and mounted Hamamatsu Orca Flash 4.0 camera. We then analyzed the videos using BEMOVI [38, 41], which returns information on the cell density, morphological traits (longest and shortest cell axis length) and movement metrics (gross speed and net speed of cells, as well as turning angle distribution). The video analysis script, including used parameter values, can be found in the Supporting Information.

### Data analysis

All statistical analyses were done using the R statistical software (version 3.5.1).

#### Predator trait space

To visualize whether the full set of trait data displayed structure depending on the evolutionary history of the predator and prey species, t-distributed stochastic neighbor embedding (t-SNE) was performed for each prey species separately using the Rtsne package [42] with a perplexity parameter of 3 owing to small sample size.

#### Beverton-Holt model fit

For analyzing the population growth dynamics of the ciliates, we implemented the Beverton-Holt population growth model [43] (Figure S3) using a Bayesian framework in Rstan, following methods used by (Fronhofer 2018; Rosenbaum et al. 2019). This function has the form of:

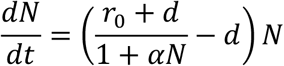

With *r_0_* being the intrinsic rate of increase, *α* the intraspecific competitive ability and *d* being the death rate in the population. Model code for fitting this function can be found on a Github repository (doi: 10.5281/zenodo.2658131). For fitting this model, we needed to provide prior information for *r_0_, d* and equilibrium density *K*. The intraspecific competitive ability *α* was later derived from the other parameter values as:

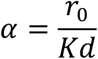

The priors (lognormal distribution) of the model were chosen in such a way that mean estimates lay close to the overall observed means, but were broad enough so the model was not constrained too strongly.

- Equilibrium population density *K*: *l*n(*K*) *~normal(9.21, 0.5)*
- Intrinsic rate of increase r0: *l*n(r0*) ~normal(−2.3, 0.5)*
- Rate of mortality d: *l*n(*d*) *~normal(−2.3, 0.5)*

Models were run with a warmup of 2,000 iterations and a chain length of 8,000 iterations.

#### Life history trait analysis

We analyzed the estimates of the life history traits obtained from the Beverton-Holt model fit (*r_0_, α*, and *K*) using linear models and model selection. We first constructed a full model with life history traits being a function of bacterial evolutionary history (evolved/ancestor), ciliate evolutionary history (evolved/ancestor) and bacterial species (seven species factors) in a full interaction model. Next, we used automated bidirectional model selection using the step function (stats package version 3.5.1) to find the best model. To avoid bias due to starting point, we fit the model both starting from the intercept model and the full model, and if model selection resulted in different models, we used AICc comparison (MuMin R-package, version 1.42.1) to select the model with the smallest AICc value.

#### Morphological and behavioral trait analysis

Morphological and behavioral data was available for every time point during the growth curve, and since we know these traits can be plastically strongly affected by density [44, 45], we had to take density into account in the model. We hence separated the analysis into two steps: first, we identified key points in the growth curves (early phase, mid-log phase and equilibrium density phase) and analysed the traits for these particular points. Secondly, we fit models over all data, but taking bacterial (using flow cytometry data) and ciliate densities into account as covariates in the statistical analysis.

We defined the early phase as the second time point in the time series, equilibrium density phase as the first time point where density was larger than 99 % of *K*, or alternatively the highest density, and the mid-log phase as the point between the early and equilibrium density phase where density was closest to 50 % of *K*. We then created statistical models for the traits (major cell axis size, gross speed of cells and turning angle distribution) as a function of bacterial evolutionary history (evolved/ancestor), ciliate evolutionary history (evolved/ancestor) and bacterial species (seven species factors) including a full interaction for the data at the particular time point. Next, we used automated bidirectional model selection to find the best fitting model. This was done separately for all three phases (early, mid-log and equilibrium density phase). We again performed model selection both starting from the intercept model and full model, and compared the 2 models using AICc comparison to identify the best model.

We then created models using all the data, where we fit major cell axis size, gross speed and turning angle distribution as a function of bacterial evolutionary history (evolved/ancestor), ciliate evolutionary history (evolved/ancestor) and bacterial species (seven species factors), ciliate population density (ln-transformed, continuous) and bacterial population density (ln-transformed, continuous), including a full interaction. For turning angle, we also did a log10 transformation of the turning angle distributions, as fitting the model on untransformed data leads to a strong deviation on the qqplot. Next, we used automated bidirectional model selection using the step function starting from intercept model and full model, and compared the 2 models using AICc comparison to select the best model.

## Results

The t-SNE maps (Figure 1) showed that the evolutionary history of the predator and prey species frequently resulted in predator divergence in trait space. Importantly, this divergence evolved from a single ancestral predator population, which was subjected to co-culture with different prey species. Prey evolution drove changes in the life history traits of the predator, including intrinsic rate of increase (*r_0_*), equilibrium density (*K*) and competitive ability (α), although the presence and strength of the effect depended on the bacterial species (ANOVA for LM on *r_0_*: prey evolution *F*_1,78_ = 15.32, *p* < 0.001; prey evolution × prey species *F*_6,78_ = 9.03, *p* < 0.001; *K*: prey evolution *F*_1,80_ = 2.43, *p* = 0.123, prey evolution × prey species *F*_6,80_ = 13.7, *p* < 0.001; α: prey evolution *F*_1,78_ = 4.79, *p* =0.031, prey evolution × prey species *F*_6,78_ = 5.40, *p* < 0.001; for full results, see Supplementary Tables S1–S3; Tables S7-S9; Figure 2). As shown in Table 2, intrinsic rate of increase of ciliates is generally lower in presence of evolved bacterial prey compared to ancestral prey, with the notable exception of *Serratia*, where intrinsic rate of increase was higher in presence of evolved prey. Also note that for three species (*Janthinobacterium, Brevundimonas* and *Pseudomonas*), evolved predators had a higher intrinsic rate of increase on evolved prey compared to ancestral prey. Changes in population equilibrium density were highly dependent on species, with four species (*Janthinobacterium*, *Brevundimonas*, *Commonas* and *Serratia*) showing higher population equilibrium density in presence of evolved prey compared to ancestral prey, and the remaining three (*Escherichia*, *Sphingomonas* and *Pseudomonas*) showing decreased population equilibrium density in presence of evolved prey compared to ancestral prey. Competitive ability typically decreased in presence of evolved prey compared to ancestral prey, with the exception of *Pseudomonas*, where competitive ability was higher in presence of evolved bacteria compared to ancestral bacteria. Also note that for *Escherichia*, *Janthinobacterium* and *Serratia* competitive ability of evolved predators was higher in presence of evolved prey compared to ancestral prey.

**Figure 1.**
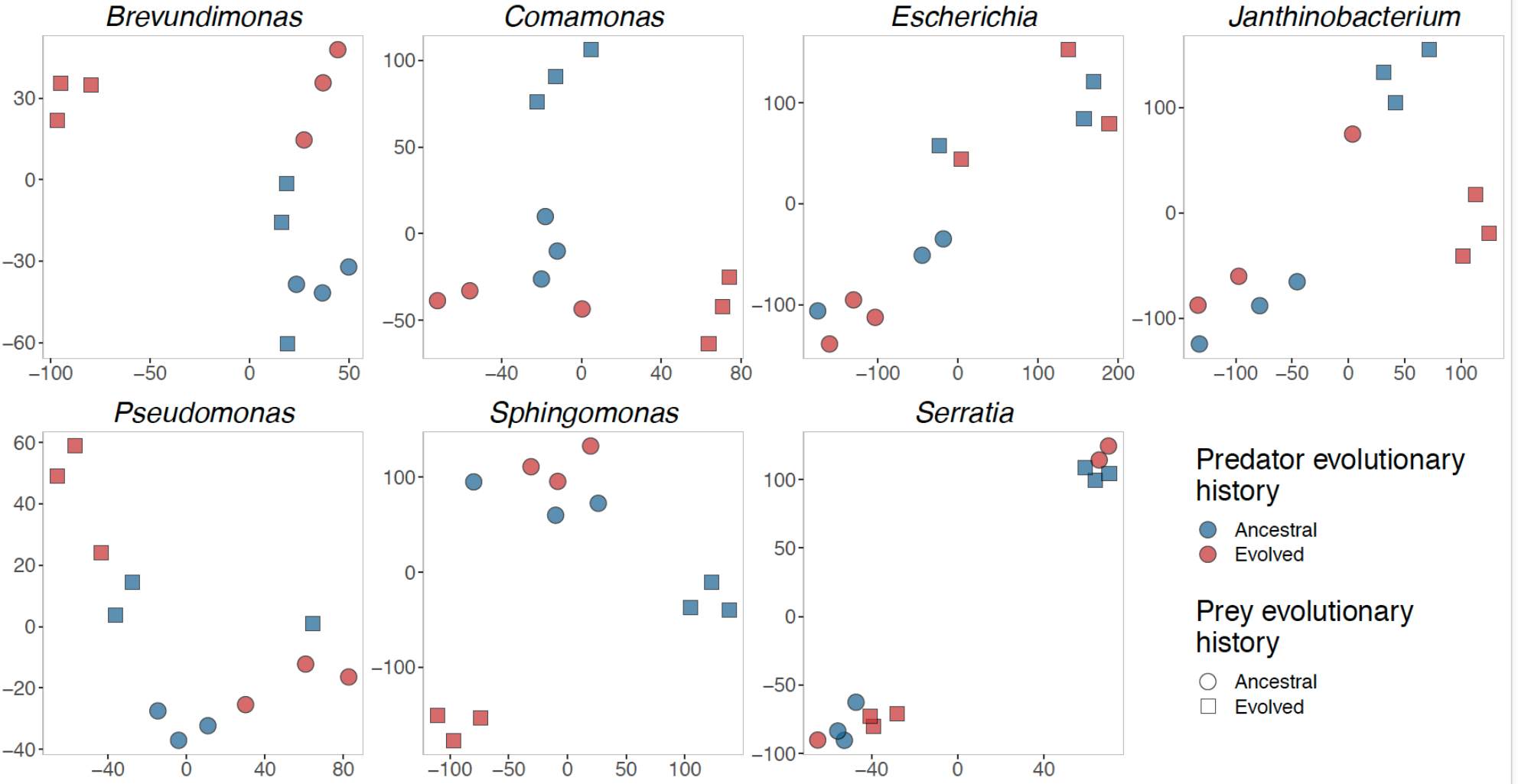
t-SNE map of contribution of predator and prey evolutionary history to predator divergence in trait space. The traits included in the analysis encompass life history (intrinsic growth rate, equilibrium density and competitive ability), morphology (cell size and biovolume) and behavior (speed and cell turning angle distribution).

**Figure 2.**
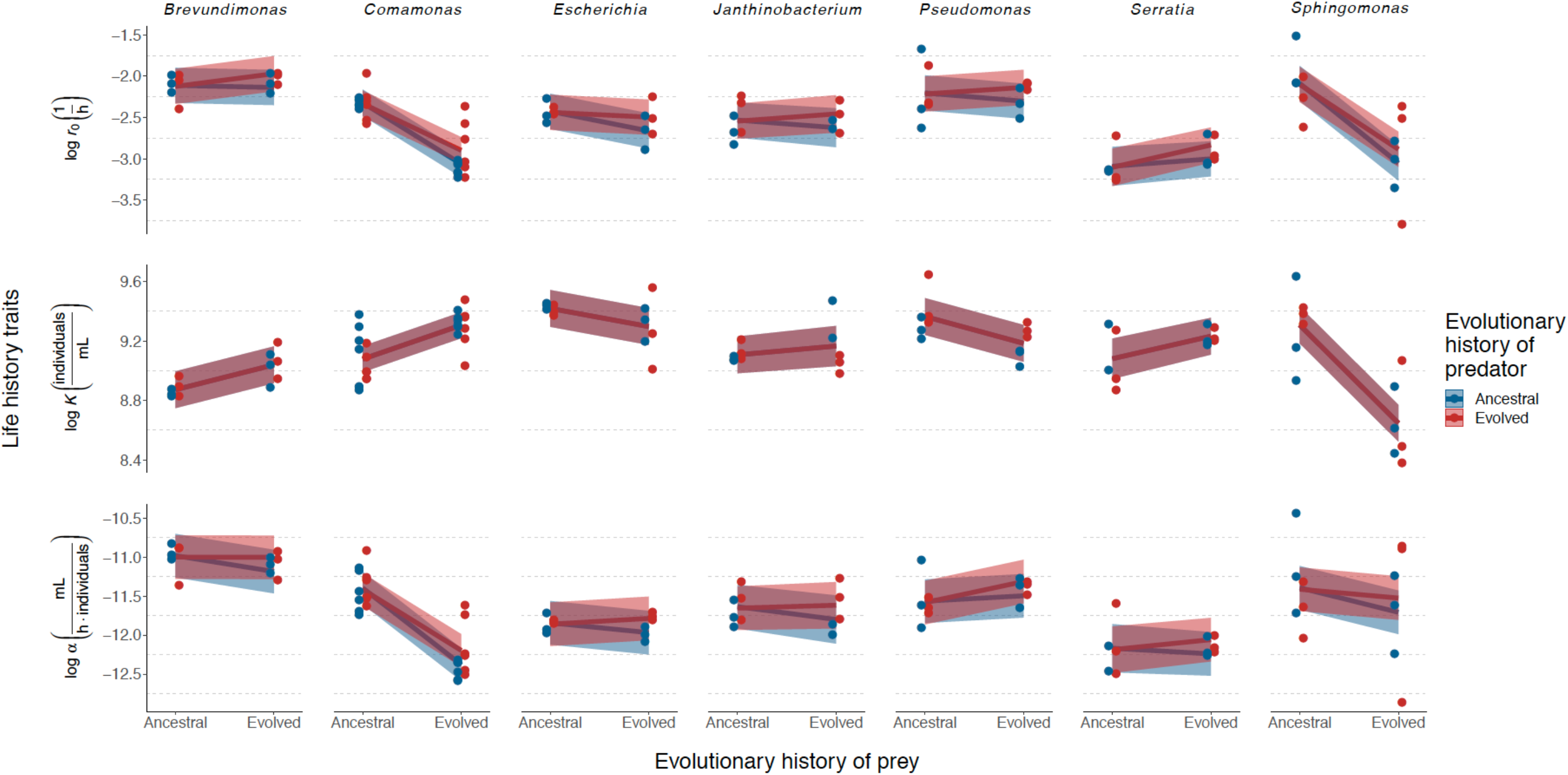
Reaction norms showing effect of evolving predator-prey interaction on life-history traits of predator (data points with linear model estimate ± 95 % confidence intervals.; N = 3 except 6 for *Comamonas*). The life-history traits for predators are parameters of Beverton-Holt continuous-time population models fitted to data, and include intrinsic growth rate (*r_0_*), equilibrium density (*K*) and competitive ability (α). The reaction norms for predators (one strain of the ciliate *Tetrahymena thermophila*) feeding on ancestral or evolved prey (seven bacterial strains indicated by genus name) are depicted separately for ancestral and evolved predators (color coding). Predators evolved with a particular prey taxon have always been coupled with ancestral or evolved populations of the same taxon, while the ancestral predator is the same for all prey taxa.

**Table 2.** Predicted change in intrinsic rate of growth (*r_0_*), population equilibrium density (*K*) and competitive ability (*α*) in presence of evolved bacteria compared to ancestral bacteria according to the linear models. The *r_0_*, *K*, and *α*-ratios are calculated as the predicted trait value (*r_0_*, *K*, or *α*) in presence of evolved bacteria divided by the predicted trait value in presence of ancestral bacteria. Note that for the *K*-ratio, since Predator evolution is excluded during model selection, predictions for ancestral and evolved predators are identical.

| Prey species | Predator evolution | $r_0$-ratio | $K$-ratio | α-ratio |
| --- | --- | --- | --- | --- |
| *E. coli* | Ancestor | 0.788 | 0.885 | 0.881 |
| *E. coli* | Evolved | 0.943 | 0.885 | 1.08 |
| *J. lividum* | Ancestor | 0.912 | 1.06 | 0.849 |
| *J. lividum* | Evolved | 1.09 | 1.06 | 1.04 |
| *S. capsulata* | Ancestor | 0.381 | 0.517 | 0.730 |
| *S. capsulata* | Evolved | 0.457 | 0.517 | 0.893 |
| *B. diminuta* | Ancestor | 0.974 | 1.18 | 0.815 |
| *B. diminuta* | Evolved | 1.17 | 1.18 | 0.997 |
| *P. fluorescens* | Ancestor | 0.904 | 0.835 | 1.07 |
| *P. fluorescens* | Evolved | 1.08 | 0.835 | 1.31 |
| *C. testosteroni* | Ancestor | 0.475 | 1.26 | 0.374 |
| *C. testosteroni* | Evolved | 0.569 | 1.26 | 0.457 |
| *S. marcescens* | Ancestor | 1.09 | 1.16 | 0.930 |
| *S. marcescens* | Evolved | 1.31 | 1.16 | 1.14 |

In contrast to life history traits, which were affected by prey evolution alone, morphological and behavioral traits of the predator were affected by predator evolution (Figure 3). However, the effect size of predator evolution was also strongly dependent on predator density (for the movement metrics, gross speed and turning angles) or both predator and prey density (cell size). Evolved predators were slightly but significantly larger than unevolved predators (ANOVA for LM on cell size: predator evolution *F*_1,767_ = 7.87, *p* = 0.005). Although there was a just significant effect indicating that this was modulated by the evolutionary history of the prey (ANOVA for LM on cell size: prey evolution *F*_1,767_ = 4.85, *p* = 0.033), the associated effect size was much smaller than predator evolution. The effect of predator evolution also depended strongly on prey densities (ANOVA for LM on cell size: log prey density) × predator evolution *F*_1,767_ = 6.87, *p* = 0.009), such that the strongest differences in cell size between ancestral and evolved predators were observed at low prey densities (cell sizes 1.2–1.3 times larger for evolved compared to ancestral ciliates; Figure 3) whereas the effects were negligible at high prey densities (approximately equal size for evolved and ancestral ciliates; for full results, see Supplementary Tables S4 and S10 and Figures S4–S6).

**Figure 3.**
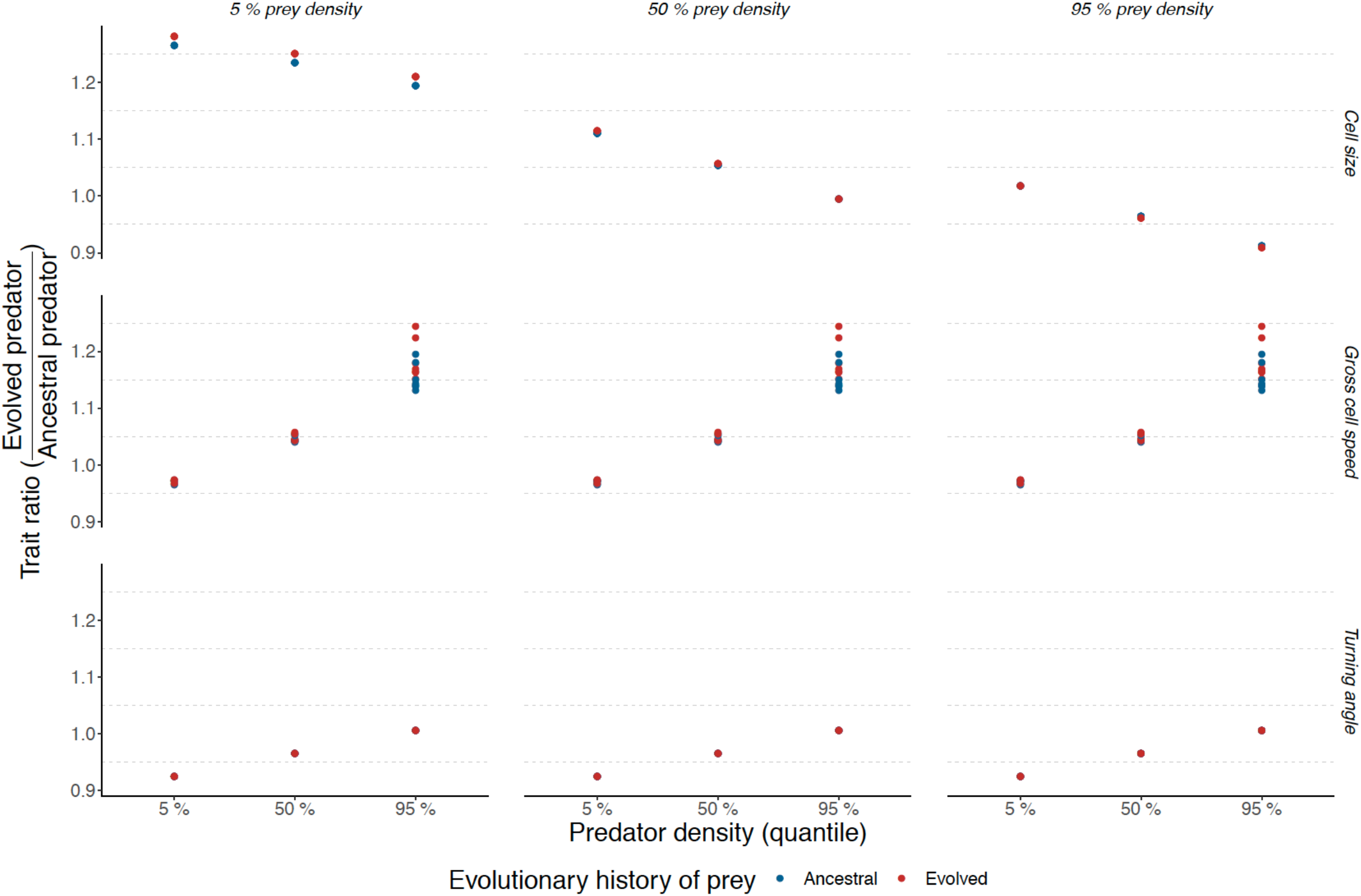
Ratios of the predicted trait values of the linear models (cell size, gross cell speed and turning angles) for the evolved predator divided by the ancestral predator at different prey densities (5 %, 50 % and 95 % quantiles) and predator densities (5 %, 50 % and 95 % quantiles). Ratios represent how ciliate traits differ between evolved and ancestral ciliates, with values of one meaning evolved and ancestral ciliates are identical, values larger than one meaning higher trait values for evolved strains, and values smaller than one higher trait values for ancestral ciliates. Note how for cell size and turning angle predictions do not differ for different prey species while for cell speed, a large variation is observed for ciliates subjected to different prey species, but only at higher predator densities. Also note the strong plastic effects on all traits associated with prey and predator densities.

The gross movement speed of predators depended on the interplay between predator density and predator or prey evolutionary history. Although evolved predators had, on average, up to 1.25 times higher speed compared to ancestral predators, this effect occurred for evolved predators at high predator densities, whereas at low predator densities, movement speed was approximately similar for ancestral and evolved ciliates (ANOVA for LM on gross speed: predator density *F*_1,763_ = 116.20, *p* < 0.001; predator evolution *F*_1,763_ = 1.90, *p* = 0.239; predator evolution × predator density *F*_1,763_ = 4.36, *p* = 0.037; Figure 3). This effect was partially counteracted by prey evolution by driving speed to a lower rate at increasing predator densities (ANOVA for LM on gross speed: prey evolution *F*_1,763_ = 2.17, *p* = 0.141; prey evolution × predator density *F*_1,763_ = 5.46, *p* = 0.020). The movement speed of ciliate cells was also dependent on identity of the prey species, with ciliates moving slower when subjected to three particular prey species *(Janthinobacterium, Pseudomonas* and *Serratia;* ANOVA for LM on gross speed: prey evolution *F*_6,763_ = 9.11, *p* < 0.001; for full results, see Tables S5 and S11 and Figure S7). Finally, predator evolution altered cell turning angle distribution, which is a proxy for the directionality of cell movement (ANOVA for LM on cell turning angle distribution: predator evolution *F*_1,56_ = 10.15, *p* = 0.001), across prey species such that evolved predator lines moved in straighter trajectories, up to a minimum of 0.92 times the turning angle distribution compared to ancestral predators. This effect was again highly dependent on predator population size (ANOVA for LM on cell turning angle distribution: predator density *F*_1,763_ = 33.90, *p* < 0.001; predator evolution × predator density *F*_1,763_ = 5.44, *p* = 0.02) with evolved predators turning approximately 0.92 as much as ancestral predators at low predator density, but turning equally much at high predator density (Figure 3). The effect of predator population size was also dependent on prey species size (ANOVA for LM on cell turning angle distribution: predator density × prey species *F*_1,763_ = 6.76, *p* < 0.001), where particularly for three prey species (*Janthinobacterium*, *Pseudomonas* and *Serratia*; for full results, see Supplementary Tables S6 and S12 and Figures S8–S10), evolved predators were more prone to continue turning less at higher densities. Note that these were the exact same prey species that had an effect on the movement speed behavior of ciliates.

## Discussion

We quantified the contribution of predator and prey evolution to predator trait change across seven different prey species in a 20-month (~600 predator generations) co-culture experiment. We expected rapid evolution of anti-predatory defense in the prey to cause impairment of predator growth [7, 46]. We expected predator evolution to be weaker in line with the life-dinner principle positing that the prey experiences stronger selection pressure since its survival (life) directly depends on defense while the predator can afford a certain measure of unsuccessful prey encounters (dinner postponement) [8, 47]. Asymmetric selection can result in dynamics other than classic arms race dynamics such as frequency-dependent cycling of traits [5], which have also been observed in microbial predator-prey systems [21]. Nevertheless, instead of escalation where predators alone impose selection pressure, we expected to also observe predator evolution, since co-evolution has been demonstrated to occur in bacteria-ciliate systems [7, 23, 46].

Prey evolution led to hypothesized changes in predator life history traits, decreasing intrinsic growth rate, equilibrium density and competitive ability, while not affecting morphological or behavioral traits in the predator. Interestingly, the strength of the effect and the life history trait affected depended on the prey species. These results may be influenced by different growth dynamics (Figures S1 and S2), defense levels or defense mechanisms [12] of the different prey species. Remarkably, against our expectation, we did not find detectable levels of adaptation in predator life history traits when prey-evolved predators fed on their respective ancestral prey species. This could be indicative of asymmetry of selection [5, 21] such that predators experience weaker selection pressure compared to prey. Slow evolutionary change for ciliate predators could also result from smaller population size (in the order of 10^4^ mL^−1^ for ciliates compared with 10^8^ mL^−1^ for bacteria), larger genome size (>100 Mb for *T. thermophila* [48] compared to <10 Mb for bacteria) or more complex genomic architecture limiting adaptive mutation supply compared to the bacterial prey. Alternatively, since improved predator growth on ancestral prey has been observed in shorter-term experiments [7, 23], growth-offense tradeoffs exacerbating over time may impose constraints on life history trait adaptation in coevolving systems in the long term [46]. It is also possible that counter-adaptations to evolved defense mechanisms, such as cell aggregation [20, 21], fail to confer an advantage to the predator when feeding on the ancestral prey.

Despite a lack of adaptation in life-history traits, evolved predators displayed both behavioral and morphological changes. Namely, increased swimming speed and body size were observed for evolved predators with certain prey species. Increased swimming speed may increase prey search efficiency [17, 22]. The role of increased body size is less clear but has been linked to the same trait since swimming speed can be a function of body size [17, 22]. Notably, predators evolved to swim in straighter trajectories across prey species, also hypothesized to increase prey search efficiency [17, 49].

Our findings have implications for interpreting data from (co-)evolving predator-prey systems. First, the pronounced impairment of predator growth traits upon prey evolution together with the lack of clear improvements in the ability of evolved predators to feed on ancestral prey types corroborate the asymmetric selection hypothesis. Second, the occurrence of predator evolution in other key traits for predator-prey interaction despite this suggests that tracking ecological changes alone may result in an underestimation of predator evolution (see also [50]). Further studies are required to identify the factors producing this constraint, such as offense-growth tradeoffs or the specificity of the advantage of improved offense to defended prey types.

## Data Availability

All code and pre-processed data needed to reproduce the ecological and evolutionary analyses will be available via Dryad.

## Author Contributions

Designed evolutionary experiment: J.C. and T.H. Performed and managed experiment: J.C. Designed physiological measurements: F.M., E.A.F., and F.A. Performed physiological measurements: F.M., E.A.F., and F.A. Analyzed data: F.M. and J.C. Wrote manuscript draft: J.C. All authors interpreted results and participated in improving the manuscript.

## Acknowledgements

We thank Veera Partanen for technical help with maintaining the evolutionary experiment and reviving samples for the physiological measurements. We thank Samuel Hürlemann for help during the lab work. This is publication ISEM-YYYY-XXX of the Institut des Sciences de l’Evolution – Montpellier. This work was funded by Academy of Finland to T.H. (project no. 106993), the University Research Priority Program (URPP) ‘Evolution in Action’ of the University of Zurich to F.A. and F.M., and Jenny and Antti Wihuri Foundation (grant no. 00190040) to J.C.

