## Supplementary Information for "Evolution in interacting species alters predator life history traits, behavior and morphology in experimental microbial communities"

**This PDF file includes:**

Supplementary methods:

R-code used in video analysis step for protist density measurements

Tables S1 to S14

Figures S1 to S14

**Supplementary methods**

*R-code (including used parameter values) used in video analysis step for protist density measurements*

```
########################################################################
# R script for analysing video files with BEMOVI (www.bemovi.info)
#
# Emanuel A. Fronhofer
#
# October 2016
########################################################################
rm(list=ls())

# load package
#library(devtools)
#install_github("efronhofer/bemovi", ref="experimental")
library(bemovi)

########################################################################
# VIDEO PARAMETERS

# video frame rate (in frames per second)
fps <- 25
# length of video (in frames)
total_frames <- 500

# measured volume (in microliter)
measured_volume <- 34.4 # for Leica M205 C with 1.6 fold magnification, sample height 0.5 mm and Hamamatsu Orca Flash 4

# size of a pixel (in micrometer)
pixel_to_scale <- 4.05 # for Leica M205 C with 1.6 fold magnification, sample height 0.5 mm and Hamamatsu Orca Flash 4
```

```
# setup
difference.lag <- 10
thresholds <- c(10,255) # don't change the second value

#######################################################################
# FILTERING PARAMETERS
#(optimized for Leica M205 C with 1.6 fold magnification, sample height 0.5 mm and Hamamatsu Orca Flash 4)
# tested species: Tet, Col, Pau, Eug, Chi, Ble, Ceph, Lox, Spi

# min and max size: area in pixels
particle_min_size <- 5
particle_max_size <- 1000

# number of adjacent frames to be considered for linking particles
trajectory_link_range <- 3
# maximum distance a particle can move between two frames
trajectory_displacement <- 16

# these values are in the units defined by the parameters above: fps (seconds), measured_volume (microliters) and
pixel_to_scale (micometers)
filter_min_net_disp <- 25
filter_min_duration <- 1
filter_detection_freq <- 0.1
filter_median_step_length <- 3
#######################################################################
# MORE PARAMETERS (USUALLY NOT CHANGED)

# UNIX
# set paths to ImageJ and particle linker standalone
IJ.path <- "/home/felix/bin/ImageJ"
to.particlelinker <- "/home/felix/bin/ParticleLinker"

# directories and file names
to.data <- paste(getwd(),"/",sep="")
```

```
video.description.folder <- "0_video_description/"
video.description.file <- "video_description.txt"
raw.video.folder <- "1_raw/"
particle.data.folder <- "2_particle_data/"
trajectory.data.folder <- "3_trajectory_data/"
temp.overlay.folder <- "4a_temp_overlays/"
overlay.folder <- "4_overlays/"
merged.data.folder <- "5_merged_data/"
ijmacs.folder <- "ijmacs/"

# RAM allocation
memory.alloc <- c(60000) # hp machine

# RAM per particle linker instance
memory.alloc.perLinker <- c(10000)
######################################################################

######################################################################
# VIDEO ANALYSIS

# identify particles
locate_and_measure_particles(to.data, raw.video.folder, particle.data.folder, difference.lag, thresholds, min_size = particle_min_size, max_size = particle_max_size, IJ.path, memory.alloc)

# link the particles
link_particles(to.data, particle.data.folder, trajectory.data.folder, linkrange = trajectory_link_range, disp = trajectory_displacement, start_vid = 1, memory = memory.alloc, memory_per_linkerProcess = memory.alloc.perLinker)

# merge info from description file and data
merge_data(to.data, particle.data.folder, trajectory.data.folder, video.description.folder, video.description.file, merged.data.folder)

# load the merged data
```

```
load(paste0(to.data, merged.data.folder, "Master.RData"))

# filter data: minimum net displacement, their duration, the detection frequency and the median step length
trajectory.data.filtered <- filter_data(trajectory.data, filter_min_net_disp, filter_min_duration, filter_detection_freq, filter_median_step_length)

# summarize trajectory data to individual-based data
morph_mvt <- summarize_trajectories(trajectory.data.filtered, calculate.median=F, write = T, to.data, merged.data.folder)

# get sample level info
summarize_populations(trajectory.data.filtered, morph_mvt, write=T, to.data, merged.data.folder, video.description.folder, video.description.file, total_frames)

# create overlays for validation
create_overlays(trajectory.data.filtered, to.data, merged.data.folder, raw.video.folder, temp.overlay.folder, overlay.folder, 2048, 2048, difference.lag, type = "label", predict_spec = F, IJ.path, contrast.enhancement = 1, memory = memory.alloc)
```

**Table S1.** ANOVA table (Type III test) for linear model on log-transformed intrinsic growth rate ($r_0$) of ciliate.

| Model terms | *d.f.* | *SS* | *F* | *p* |
|---|---|---|---|---|
| Prey evolution | 1 | 0.93 | 15.32 | **< 0.001** |
| Predator evolution | 1 | 0.14 | 2.30 | 0.134 |
| Prey species | 6 | 6.41 | 17.51 | **< 0.001** |
| Prey evolution × predator evolution | 1 | 0.19 | 3.10 | 0.082 |
| Prey evolution × prey species | 6 | 3.31 | 9.03 | **< 0.001** |
| Residuals | 78 | 4.7589 | | |

**Table S2.** ANOVA table (Type III test) for linear model on log-transformed competitive ability (α) of ciliate.

| Model terms | *d.f.* | *SS* | *F* | *p* |
|---|---|---|---|---|
| Prey evolution | 1 | 0.5 | 4.7927 | **0.031** |
| Prey species | 6 | 9.3 | 14.4450 | **< 0.001** |
| Prey evolution × prey species | 6 | 3.5 | 5.3986 | **< 0.001** |
| Residuals | 80 | 8.6 | | |

**Table S3.** ANOVA table (Type III test) for linear model on log-transformed equilibrium density ($K$) of ciliate.

| Model terms | *d.f.* | *SS* | *F* | *p* |
|---|---|---|---|---|
| Prey evolution | 1 | 0.1 | 2.43 | 0.123 |
| Prey species | 6 | 1.6 | 11.2 | **< 0.001** |
| Prey evolution × prey species | 6 | 1.9 | 13.7 | **< 0.001** |
| Residuals | 80 | 1.9 | | |

**Table S4.** ANOVA table (Type III test) for linear model on cell size of ciliate.

| Model terms | *d.f.* | *SS* | *F* | *p* |
|---|---|---|---|---|
| Prey evolution | 1 | 483 | 4.58 | **0.033** |
| Predator evolution | 1 | 830 | 7.87 | **0.005** |
| Log prey population size | 1 | 24 | 0.23 | 0.633 |
| Log predator population size | 1 | 722 | 6.85 | **0.009** |
| Log prey population size × predator evolution | 1 | 725 | 6.87 | **0.009** |
| Log predator population size × predator evolution | 1 | 266 | 2.52 | 0.113 |
| Log predator population size × log prey population size | 1 | 725 | 5.24 | **0.022** |
| Residuals | 767 | 80886 | | |

**Table S5.** ANOVA table (Type III test) for linear model on gross speed of ciliate.

| Model terms | *d.f.* | *SS* | *F* | *p* |
|---|---|---|---|---|
| Prey evolution | 1 | 14526 | 2.17 | 0.141 |
| Predator evolution | 1 | 9314 | 1.90 | 0.239 |
| Prey species | 6 | 366264 | 9.11 | **< 0.001** |
| Log predator population size | 1 | 778688 | 116.20 | **< 0.001** |
| Log predator population size × prey evolution | 1 | 36601 | 5.46 | **0.020** |
| Log predator population size × predator evolution | 1 | 29248 | 4.36 | **0.037** |
| Residuals | 763 | 5113281 | | |

**Table S6.** ANOVA table (Type III test) for linear model on cell turning angle distribution of ciliate.

| Model terms | *d.f.* | *SS* | *F* | *p* |
|---|---|---|---|---|
| Predator evolution | 1 | 0.27 | 10.15 | **0.001** |
| Prey evolution | 1 | 0.08 | 3.10 | 0.079 |
| Prey species | 6 | 1.98 | 12.40 | **<0.001** |
| Log prey population size | 1 | 0.27 | 10.26 | **0.001** |
| Log predator population size | 1 | 0.90 | 33.90 | **< 0.001** |
| Log predator population size × predator evolution | 1 | 0.14 | 5.44 | **0.02** |
| Log predator population size × prey species | 1 | 1.08 | 6.76 | **< 0.001** |
| Log prey population size × prey evolution | 1 | 0.07 | 2.76 | 0.097 |
| Residuals | 756 | 20.09 | | |

**Table S7.** Summary table for linear model on log-transformed intrinsic growth rate ($r_0$) of ciliate.

| *Model terms* | *Estimate* | *SE* | *t-value* | *Pr(>\|t\|)* | *Significance* |
|---|---|---|---|---|---|
| Intercept (*B. diminuta)* | -2.111 | 0.107 | -19.707 | <0.001 | *** |
| Prey evolution (evolved) | -0.026 | 0.151 | -0.172 | 0.864 | |
| Predator evolution (evolved) | -0.013 | 0.072 | -0.174 | 0.862 | |
| Prey species (*C. testosteroni*) | -0.210 | 0.124 | -1.697 | 0.094 | . |
| Prey species (*E. coli*) | -0.314 | 0.143 | -2.201 | 0.031 | * |
| Prey species (*J. lividum*) | -0.421 | 0.143 | -2.949 | 0.004 | ** |
| Prey species (*P. fluorescens*) | -0.092 | 0.143 | -0.642 | 0.523 | |
| Prey species (*S. capsulata*) | 0.023 | 0.143 | 0.160 | 0.873 | |
| Prey species (*S. marcescens*) | -0.980 | 0.150 | -6.542 | <0.001 | *** |
| Prey evolution (evolved) * Predater evolution (evolved) | 0.180 | 0.102 | 1.762 | 0.082 | . |
| Prey evolution (evolved) * Prey species (*C. testosteroni*) | -0.718 | 0.175 | -4.112 | <0.001 | *** |
| Prey evolution (evolved) * Prey species (*E. coli*) | -0.212 | 0.202 | -1.052 | 0.296 | |
| Prey evolution (evolved) * Prey species (*J. lividum*) | -0.066 | 0.207 | -0.321 | 0.749 | |
| Prey evolution (evolved) * Prey species (*P. fluorescens*) | -0.075 | 0.202 | -0.370 | 0.713 | |
| Prey evolution (evolved) * Prey species (*S. capsulata*) | -0.938 | 0.202 | -4.649 | <0.001 | *** |
| Prey evolution (evolved) * Prey species (*S. marcescens*) | 0.116 | 0.207 | 0.562 | 0.576 | |

**Table S8.** Summary table for linear model on log-transformed competitive ability ($\alpha$) of ciliate.

| *Model terms* | *Estimate* | *SE* | *t-value* | *Pr(>\|t\|)* | *Significance* |
|---|---|---|---|---|---|
| Intercept (*B. diminuta)* | -10.980 | 0.141 | -78.121 | <0.001 | *** |
| Prey evolution (evolved) | -0.204 | 0.199 | -1.028 | 0.307 | |
| Predator evolution (evolved) | -0.018 | 0.095 | -0.195 | 0.846 | |
| Prey species (*C. testosteroni*) | -0.416 | 0.162 | -2.565 | 0.012 | * |
| Prey species (*E. coli*) | -0.861 | 0.187 | -4.602 | <0.001 | *** |
| Prey species (*J. lividum*) | -0.655 | 0.187 | -3.501 | <0.001 | *** |
| Prey species (*P. fluorescens*) | -0.584 | 0.187 | -3.123 | 0.003 | ** |
| Prey species (*S. capsulata*) | -0.412 | 0.187 | -2.203 | 0.031 | * |
| Prey species (*S. marcescens*) | -1.188 | 0.197 | -6.044 | 0.000 | *** |
| Prey evolution (evolved) * Predater evolution (evolved) | 0.201 | 0.134 | 1.502 | 0.137 | |
| Prey evolution (evolved) * Prey species (*C. testosteroni*) | -0.779 | 0.229 | -3.399 | 0.001 | ** |
| Prey evolution (evolved) * Prey species (*E. coli*) | 0.078 | 0.265 | 0.293 | 0.770 | |
| Prey evolution (evolved) * Prey species (*J. lividum*) | 0.040 | 0.271 | 0.149 | 0.882 | |
| Prey evolution (evolved) * Prey species (*P. fluorescens*) | 0.274 | 0.265 | 1.034 | 0.304 | |
| Prey evolution (evolved) * Prey species (*S. capsulata*) | -0.110 | 0.265 | -0.416 | 0.678 | |
| Prey evolution (evolved) * Prey species (*S. marcescens*) | 0.132 | 0.271 | 0.485 | 0.629 | |

**Table S9.** Summary table for linear model on log-transformed equilibrium density ($K$) of ciliate.

| *Model terms* | *Estimate* | *SE* | *t-value* | *Pr(>\|t\|)* | *Significance* |
|---|---|---|---|---|---|
| Intercept (*B. diminuta)* | 8.873 | 0.062 | 142.519 | <0.001 | *** |
| Prey evolution (evolved) | 0.168 | 0.088 | 1.904 | 0.060 | . |
| Prey species (*C. testosteroni*) | 0.206 | 0.076 | 2.703 | 0.008 | ** |
| Prey species (*E. coli*) | 0.547 | 0.088 | 6.218 | <0.001 | *** |
| Prey species (*J. lividum*) | 0.235 | 0.088 | 2.664 | 0.009 | ** |
| Prey species (*P. fluorescens*) | 0.493 | 0.088 | 5.599 | <0.001 | *** |
| Prey species (*S. capsulata*) | 0.435 | 0.088 | 4.943 | <0.001 | *** |
| Prey species (*S. marcescens*) | 0.209 | 0.092 | 2.259 | 0.027 | * |
| Prey evolution (evolved) * Prey species (*C. testosteroni*) | 0.061 | 0.108 | 0.564 | 0.574 | |
| Prey evolution (evolved) * Prey species (*E. coli*) | -0.290 | 0.125 | -2.328 | 0.022 | * |
| Prey evolution (evolved) * Prey species (*J. lividum*) | -0.108 | 0.128 | -0.848 | 0.399 | |
| Prey evolution (evolved) * Prey species (*P. fluorescens*) | -0.348 | 0.125 | -2.797 | 0.006 | ** |
| Prey evolution (evolved) * Prey species (*S. capsulata*) | -0.828 | 0.125 | -6.646 | <0.001 | *** |
| Prey evolution (evolved) * Prey species (*S. marcescens*) | -0.016 | 0.128 | -0.125 | 0.901 | |

**Table S10.** Summary table for linear model on cell size of ciliate.

| *Model terms* | *Estimate* | *SE* | *t-value* | *Pr(>\|t\|)* | *Significance* |
|---|---|---|---|---|---|
| Intercept | 11.142 | 20.710 | 0.538 | 0.591 | |
| Log (Predator density) | -5.956 | 2.431 | -2.450 | 0.015 | * |
| Log (Prey density) | 1.323 | 1.044 | 1.267 | 0.206 | |
| Predator evolution (evolved) | 39.120 | 13.945 | 2.805 | 0.005 | ** |
| Prey evolution (evolved) | -1.629 | 0.761 | -2.140 | 0.033 | * |
| Log (Prey density) * Log (Predator density) | 0.287 | 0.125 | 2.289 | 0.022 | * |
| Log (Prey density) * Predator evolution (evolved) | -1.800 | 0.687 | -2.622 | 0.009 | ** |
| Log (Predator density) * Predator evolution (evolved) | -0.509 | 0.321 | -1.588 | 0.113 | |

**Table S11.** Summary table for linear model on gross speed of ciliate.

| *Model terms* | *Estimate* | *SE* | *t-value* | *Pr(>|t|)* | *Significance* |
|---|---|---|---|---|---|
| Intercept (*B. diminuta)* | 341.529 | 15.332 | 22.276 | <0.001 | *** |
| Log (Predator density) | -12.184 | 1.922 | -6.340 | <0.001 | *** |
| Prey species (*C. testosteroni*) | -13.444 | 9.735 | -1.381 | 0.168 | |
| Prey species (*E. coli*) | -14.996 | 11.327 | -1.324 | 0.186 | |
| Prey species (*J. lividum*) | -72.371 | 12.319 | -5.875 | <0.001 | *** |
| Prey species (*P. fluorescens*) | -29.134 | 11.603 | -2.511 | 0.012 | * |
| Prey species (*S. capsulata*) | -19.143 | 11.293 | -1.695 | 0.091 | . |
| Prey species (*S. marcescens*) | -61.507 | 12.695 | -4.845 | <0.001 | *** |
| Predator evolution (evolved) | -17.989 | 15.259 | -1.179 | 0.239 | |
| Prey evolution (evolved) | 22.698 | 15.417 | 1.472 | 0.141 | |
| Log (Predator density) * Predator evolution (evolved) | 4.703 | 2.251 | 2.089 | 0.037 | * |
| Log (Predator density) * Prey evolution (evolved) | -5.316 | 2.275 | -2.337 | 0.020 | * |

**Table S12.** Summary table for linear model on cell turning angle distribution of ciliate.

| Model terms | Estimate | SE | t-value | Pr(>|t|) | Significance |
|---|---|---|---|---|---|
| Intercept (*B. diminuta)* | -0.208 | 0.139 | -1.499 | 0.134 | |
| Prey species (*C. testosteroni*) | 0.068 | 0.064 | 1.057 | 0.291 | |
| Prey species (*E. coli*) | 0.044 | 0.070 | 0.637 | 0.525 | |
| Prey species (*J. lividum*) | 0.379 | 0.068 | 5.599 | <0.001 | *** |
| Prey species (*P. fluorescens*) | 0.304 | 0.066 | 4.597 | <0.001 | *** |
| Prey species (*S. capsulata*) | 0.075 | 0.073 | 1.028 | 0.304 | |
| Prey species (*S. marcescens*) | 0.128 | 0.069 | 1.849 | 0.065 | . |
| Log (Predator density) | -0.002 | 0.008 | -0.236 | 0.813 | |
| Log (Prey density) | -0.008 | 0.006 | -1.188 | 0.235 | |
| Predator evolution (evolved) | -0.100 | 0.031 | -3.187 | 0.002 | ** |
| Prey evolution (evolved) | 0.305 | 0.174 | 1.759 | 0.079 | . |
| Log (Predator density) * Prey species (*C. testosteroni*) | -0.014 | 0.009 | -1.611 | 0.108 | |
| Log (Predator density) * Prey species (*E. coli*) | -0.005 | 0.010 | -0.490 | 0.624 | |
| Log (Predator density) * Prey species (*J. lividum*) | -0.041 | 0.010 | -4.281 | <0.001 | *** |
| Log (Predator density) * Prey species (*P. fluorescens*) | -0.036 | 0.009 | -3.928 | <0.001 | *** |
| Log (Predator density) * Prey species (*S. capsulata*) | -0.006 | 0.011 | -0.601 | 0.548 | |
| Log (Predator density) * Prey species (*S. marcescens*) | -0.023 | 0.011 | -2.165 | 0.031 | * |
| Log (Predator density) * Predator evolution (evolved) | 0.011 | 0.005 | 2.333 | 0.020 | * |
| Log (Prey density) * Prey evolution (evolved) | -0.015 | 0.009 | -1.661 | 0.097 | . |

**Table S13.** Summary statistics for the Beverton-Holt model fitting. Rows represent growth curve fits for individual populations, showing mean and standard deviation for the posterior distribution of r0, α, K and d. For r0, α and d, quality statistics (effective sample size N_EFF and Rhat scores) are also shown. Note the absence of Rhat and N_eff values for α, since α is calculated from the other parameters. Large effective sample sizes and Rhat scores that are not too divergent from 1 are indicative for a healthy MCMC chain, and hence of good model convergence.

| *Name* | *log($r_0$) mean* | *log($r_0$) sd* | *log($r_0$) N_EFF* | *log($r_0$) Rhat* | *log(K) mean* | *log(K) sd* | *log(K) N_EFF* | *log(K) Rhat* | *log(α) mean* | *log(α) sd* | *log(d) mean* | *log(d) sd* | *log(d) N_EFF* | *log(d) Rhat* |
|---|---|---|---|---|---|---|---|---|---|---|---|---|---|---|
| sample_00030 | -2,566 | 0,205 | 7596 | 1,000 | 9,411 | 0,440 | 10086 | 1,000 | -11,977 | 0,491 | -2,100 | 0,936 | 9458 | 1,000 |
| sample_00031 | -2,271 | 0,196 | 5751 | 1,000 | 9,444 | 0,403 | 8138 | 1,000 | -11,716 | 0,456 | -2,017 | 0,927 | 7158 | 1,000 |
| sample_00032 | -2,476 | 0,188 | 4391 | 1,000 | 9,455 | 0,389 | 8546 | 1,000 | -11,932 | 0,445 | -2,023 | 0,938 | 8735 | 1,000 |
| sample_00033 | -2,825 | 0,279 | 9101 | 1,001 | 9,073 | 0,481 | 12073 | 1,000 | -11,898 | 0,577 | -2,332 | 1,015 | 10082 | 1,000 |
| sample_00034 | -2,483 | 0,217 | 4885 | 1,000 | 9,069 | 0,419 | 9973 | 1,000 | -11,552 | 0,475 | -2,167 | 0,974 | 9104 | 1,000 |
| sample_00035 | -2,675 | 0,292 | 7540 | 1,000 | 9,100 | 0,468 | 11232 | 1,000 | -11,775 | 0,578 | -2,307 | 0,999 | 11251 | 1,000 |
| sample_00036 | -1,509 | 0,196 | 3955 | 1,000 | 8,933 | 0,277 | 6211 | 1,000 | -10,442 | 0,364 | -1,836 | 0,878 | 6888 | 1,000 |
| sample_00037 | -2,086 | 0,127 | 4082 | 1,001 | 9,158 | 0,220 | 5726 | 1,000 | -11,245 | 0,266 | -1,747 | 0,865 | 4913 | 1,000 |
| sample_00038 | -2,080 | 0,149 | 4901 | 1,000 | 9,638 | 0,269 | 6809 | 1,000 | -11,718 | 0,327 | -1,766 | 0,839 | 5922 | 1,000 |
| sample_00039 | -1,982 | 0,225 | 6460 | 1,000 | 8,840 | 0,356 | 8636 | 1,000 | -10,822 | 0,450 | -2,010 | 0,915 | 8449 | 1,000 |
| sample_00040 | -2,092 | 0,244 | 5798 | 1,000 | 8,878 | 0,379 | 8892 | 1,000 | -10,971 | 0,474 | -2,026 | 0,912 | 7816 | 1,000 |
| sample_00041 | -2,195 | 0,257 | 6000 | 1,000 | 8,828 | 0,388 | 7360 | 1,000 | -11,023 | 0,489 | -2,139 | 0,950 | 7739 | 1,000 |
| sample_00042 | -1,673 | 0,239 | 3584 | 1,000 | 9,362 | 0,370 | 7318 | 1,000 | -11,035 | 0,434 | -1,910 | 0,871 | 6303 | 1,000 |
| sample_00043 | -2,632 | 0,292 | 8438 | 1,000 | 9,274 | 0,466 | 10593 | 1,000 | -11,907 | 0,569 | -2,256 | 0,984 | 11079 | 1,000 |
| sample_00044 | -2,399 | 0,203 | 7208 | 1,000 | 9,217 | 0,412 | 10505 | 1,000 | -11,616 | 0,469 | -2,083 | 0,940 | 10133 | 1,000 |
| sample_00045 | -2,398 | 0,231 | 6805 | 1,000 | 9,297 | 0,368 | 9565 | 1,000 | -11,695 | 0,460 | -2,253 | 0,956 | 8551 | 1,000 |
| sample_00046 | -2,296 | 0,198 | 5070 | 1,000 | 9,145 | 0,348 | 6422 | 1,000 | -11,441 | 0,418 | -2,062 | 0,929 | 6743 | 1,000 |
| sample_00047 | -2,349 | 0,222 | 4770 | 1,001 | 9,204 | 0,355 | 7649 | 1,000 | -11,553 | 0,442 | -2,111 | 0,949 | 7689 | 1,000 |
| sample_00048 | -2,264 | 0,222 | 5000 | 1,000 | 8,872 | 0,366 | 6340 | 1,000 | -11,136 | 0,440 | -2,028 | 0,961 | 7220 | 1,000 |

| | | | | | | | | | | | | | | |
|---|---|---|---|---|---|---|---|---|---|---|---|---|---|---|
| sample_00049 | -2,278 | 0,237 | 5734 | 1,000 | 8,891 | 0,358 | 7157 | 1,000 | -11,169 | 0,447 | -2,067 | 0,955 | 6838 | 1,000 |
| sample_00050 | -2,355 | 0,150 | 4181 | 1,000 | 9,380 | 0,280 | 6514 | 1,000 | -11,735 | 0,337 | -2,016 | 0,905 | 6814 | 1,000 |
| sample_00051 | NA | NA | NA | NA | NA | NA | NA | NA | NA | NA | NA | NA | NA | NA |
| sample_00052 | -3,152 | 0,242 | 5611 | 1,000 | 9,313 | 0,483 | 9476 | 1,000 | -12,465 | 0,564 | -2,232 | 0,976 | 10639 | 1,000 |
| sample_00053 | -3,136 | 0,684 | 1885 | 1,000 | 9,003 | 0,529 | 4018 | 1,000 | -12,139 | 0,970 | -2,444 | 1,083 | 5256 | 1,000 |
| sample_00054 | -2,475 | 0,182 | 5627 | 1,000 | 9,420 | 0,382 | 8121 | 1,000 | -11,896 | 0,431 | -1,989 | 0,909 | 9882 | 1,000 |
| sample_00055 | -2,650 | 0,226 | 4337 | 1,000 | 9,345 | 0,393 | 8882 | 1,000 | -11,995 | 0,481 | -2,161 | 0,957 | 7340 | 1,000 |
| sample_00056 | -2,889 | 0,413 | 8531 | 1,000 | 9,200 | 0,495 | 12297 | 1,000 | -12,089 | 0,651 | -2,308 | 1,008 | 11962 | 1,000 |
| sample_00057 | -2,639 | 0,269 | 8348 | 1,000 | 9,220 | 0,455 | 12106 | 1,000 | -11,859 | 0,543 | -2,248 | 0,971 | 11052 | 1,000 |
| sample_00058 | NA | NA | NA | NA | NA | NA | NA | NA | NA | NA | NA | NA | NA | NA |
| sample_00059 | -2,529 | 0,195 | 5614 | 1,001 | 9,470 | 0,450 | 5945 | 1,000 | -11,999 | 0,494 | -2,066 | 0,922 | 6375 | 1,000 |
| sample_00060 | -3,353 | 0,596 | 3432 | 1,000 | 8,891 | 0,575 | 6697 | 1,000 | -12,244 | 0,973 | -2,614 | 1,198 | 6851 | 1,000 |
| sample_00061 | -3,002 | 0,632 | 2524 | 1,001 | 8,612 | 0,609 | 3861 | 1,000 | -11,614 | 1,063 | -3,263 | 1,522 | 3075 | 1,001 |
| sample_00062 | -2,788 | 0,559 | 3602 | 1,000 | 8,446 | 0,600 | 3795 | 1,000 | -11,234 | 0,969 | -3,189 | 1,429 | 4934 | 1,000 |
| sample_00063 | -1,970 | 0,252 | 6403 | 1,000 | 9,040 | 0,367 | 8916 | 1,000 | -11,010 | 0,465 | -1,960 | 0,901 | 9479 | 1,000 |
| sample_00064 | -2,095 | 0,199 | 5027 | 1,000 | 9,111 | 0,272 | 7725 | 1,000 | -11,205 | 0,358 | -1,867 | 0,865 | 7731 | 1,000 |
| sample_00065 | -2,205 | 0,262 | 6197 | 1,000 | 8,888 | 0,363 | 7911 | 1,000 | -11,093 | 0,468 | -2,097 | 0,952 | 8552 | 1,000 |
| sample_00066 | -2,511 | 0,205 | 6804 | 1,000 | 9,135 | 0,431 | 9383 | 1,000 | -11,646 | 0,485 | -2,169 | 0,964 | 10682 | 1,000 |
| sample_00067 | -2,139 | 0,218 | 6529 | 1,000 | 9,127 | 0,378 | 9690 | 1,000 | -11,266 | 0,447 | -1,963 | 0,907 | 9782 | 1,000 |
| sample_00068 | -2,334 | 0,161 | 3430 | 1,001 | 9,027 | 0,381 | 8259 | 1,001 | -11,361 | 0,429 | -1,962 | 0,921 | 6484 | 1,000 |
| sample_00069 | -3,053 | 0,274 | 7379 | 1,000 | 9,298 | 0,481 | 11287 | 1,000 | -12,351 | 0,568 | -2,267 | 0,986 | 9991 | 1,000 |
| sample_00070 | -3,168 | 0,198 | 4374 | 1,000 | 9,308 | 0,459 | 8336 | 1,000 | -12,476 | 0,522 | -2,225 | 0,953 | 9085 | 1,000 |
| sample_00071 | -3,175 | 0,204 | 4892 | 1,000 | 9,407 | 0,456 | 8374 | 1,000 | -12,581 | 0,524 | -2,184 | 0,963 | 8918 | 1,000 |
| sample_00072 | -3,072 | 0,317 | 5796 | 1,000 | 9,246 | 0,498 | 9779 | 1,000 | -12,319 | 0,631 | -2,336 | 0,993 | 11166 | 1,000 |
| sample_00073 | -3,011 | 0,191 | 6272 | 1,000 | 9,337 | 0,437 | 10815 | 1,000 | -12,348 | 0,495 | -2,214 | 0,974 | 10006 | 1,000 |
| sample_00074 | -3,224 | 0,224 | 3461 | 1,000 | 9,355 | 0,438 | 6628 | 1,000 | -12,579 | 0,538 | -2,225 | 0,986 | 6706 | 1,000 |

| | | | | | | | | | | | | | | |
|---|---|---|---|---|---|---|---|---|---|---|---|---|---|---|
| sample_00075 | -3,051 | 0,390 | 6521 | 1,000 | 9,176 | 0,506 | 10493 | 1,000 | -12,227 | 0,659 | -2,368 | 1,005 | 10026 | 1,000 |
| sample_00076 | -3,070 | 0,436 | 4765 | 1,000 | 9,196 | 0,504 | 9287 | 1,000 | -12,266 | 0,680 | -2,321 | 1,027 | 8960 | 1,000 |
| sample_00077 | -2,697 | 0,267 | 5555 | 1,000 | 9,315 | 0,431 | 7367 | 1,000 | -12,012 | 0,540 | -2,211 | 0,954 | 6439 | 1,000 |
| sample_00078 | -2,378 | 0,226 | 5482 | 1,000 | 9,441 | 0,402 | 10503 | 1,001 | -11,819 | 0,476 | -2,024 | 0,909 | 9211 | 1,000 |
| sample_00079 | -2,458 | 0,218 | 7480 | 1,000 | 9,394 | 0,419 | 10267 | 1,000 | -11,852 | 0,481 | -2,086 | 0,910 | 9988 | 1,000 |
| sample_00080 | -2,436 | 0,267 | 5465 | 1,001 | 9,374 | 0,402 | 7384 | 1,000 | -11,810 | 0,507 | -2,134 | 0,943 | 8092 | 1,000 |
| sample_00081 | -2,236 | 0,245 | 5390 | 1,000 | 9,079 | 0,418 | 7465 | 1,000 | -11,315 | 0,489 | -2,125 | 0,959 | 7673 | 1,000 |
| sample_00082 | -2,685 | 0,328 | 8940 | 1,000 | 9,116 | 0,468 | 10390 | 1,000 | -11,800 | 0,584 | -2,310 | 1,025 | 10137 | 1,000 |
| sample_00083 | -2,321 | 0,254 | 6832 | 1,001 | 9,206 | 0,425 | 12528 | 1,000 | -11,528 | 0,504 | -2,104 | 0,949 | 9795 | 1,000 |
| sample_00084 | -2,263 | 0,191 | 4572 | 1,001 | 9,381 | 0,309 | 8221 | 1,001 | -11,644 | 0,383 | -2,066 | 0,914 | 8560 | 1,000 |
| sample_00085 | -2,008 | 0,174 | 5096 | 1,000 | 9,311 | 0,290 | 7351 | 1,000 | -11,320 | 0,358 | -1,994 | 0,880 | 8367 | 1,000 |
| sample_00086 | -2,618 | 0,190 | 3826 | 1,000 | 9,426 | 0,318 | 5466 | 1,000 | -12,044 | 0,414 | -2,493 | 0,966 | 5718 | 1,000 |
| sample_00087 | -2,047 | 0,279 | 8179 | 1,000 | 8,832 | 0,385 | 8669 | 1,000 | -10,880 | 0,501 | -2,082 | 0,937 | 10311 | 1,000 |
| sample_00088 | -2,399 | 0,264 | 5221 | 1,001 | 8,966 | 0,389 | 10502 | 1,000 | -11,364 | 0,494 | -2,213 | 0,967 | 10118 | 1,000 |
| sample_00089 | -1,988 | 0,262 | 7163 | 1,000 | 8,891 | 0,373 | 9422 | 1,000 | -10,879 | 0,474 | -2,004 | 0,902 | 9342 | 1,000 |
| sample_00090 | -1,872 | 0,151 | 4758 | 1,000 | 9,647 | 0,332 | 5705 | 1,000 | -11,520 | 0,364 | -1,730 | 0,850 | 6867 | 1,000 |
| sample_00091 | -2,347 | 0,260 | 5342 | 1,000 | 9,368 | 0,417 | 10538 | 1,000 | -11,715 | 0,525 | -2,117 | 0,932 | 8601 | 1,000 |
| sample_00092 | -2,327 | 0,246 | 5151 | 1,000 | 9,325 | 0,376 | 10109 | 1,000 | -11,652 | 0,473 | -2,050 | 0,922 | 9267 | 1,000 |
| sample_00093 | -2,300 | 0,208 | 4756 | 1,000 | 8,995 | 0,334 | 7793 | 1,000 | -11,295 | 0,414 | -2,023 | 0,903 | 6347 | 1,000 |
| sample_00094 | -2,264 | 0,156 | 5092 | 1,000 | 8,992 | 0,252 | 5223 | 1,000 | -11,256 | 0,315 | -1,838 | 0,898 | 5265 | 1,000 |
| sample_00095 | -2,531 | 0,248 | 5116 | 1,000 | 9,094 | 0,398 | 9186 | 1,000 | -11,625 | 0,489 | -2,194 | 0,974 | 8949 | 1,000 |
| sample_00096 | -1,969 | 0,242 | 5559 | 1,000 | 8,945 | 0,329 | 6672 | 1,000 | -10,914 | 0,428 | -1,983 | 0,918 | 6885 | 1,000 |
| sample_00097 | -2,346 | 0,331 | 8855 | 1,000 | 9,184 | 0,459 | 12195 | 1,000 | -11,530 | 0,580 | -2,289 | 1,001 | 11671 | 1,000 |
| sample_00098 | -2,571 | 0,253 | 5226 | 1,000 | 8,945 | 0,391 | 7044 | 1,000 | -11,515 | 0,491 | -2,269 | 1,006 | 9423 | 1,000 |
| sample_00099 | -3,262 | 0,449 | 3889 | 1,000 | 8,945 | 0,525 | 7750 | 1,000 | -12,207 | 0,779 | -2,577 | 1,125 | 7451 | 1,000 |
| sample_00100 | -3,221 | 0,286 | 4523 | 1,000 | 9,272 | 0,480 | 8471 | 1,000 | -12,493 | 0,603 | -2,277 | 1,008 | 8037 | 1,000 |

| | | | | | | | | | | | | | | |
|---|---|---|---|---|---|---|---|---|---|---|---|---|---|---|
| sample_00101 | -2,719 | 0,495 | 8428 | 1,000 | 8,873 | 0,522 | 8793 | 1,000 | -11,591 | 0,804 | -2,633 | 1,114 | 8919 | 1,000 |
| sample_00102 | -2,247 | 0,211 | 5219 | 1,000 | 9,562 | 0,378 | 8011 | 1,000 | -11,809 | 0,445 | -1,938 | 0,885 | 8359 | 1,000 |
| sample_00103 | -2,516 | 0,266 | 4494 | 1,001 | 9,249 | 0,374 | 7889 | 1,000 | -11,765 | 0,494 | -2,153 | 0,936 | 7723 | 1,000 |
| sample_00104 | -2,697 | 0,236 | 6083 | 1,000 | 9,011 | 0,410 | 8933 | 1,000 | -11,709 | 0,509 | -2,388 | 1,008 | 10108 | 1,000 |
| sample_00105 | -2,463 | 0,202 | 5443 | 1,000 | 9,059 | 0,399 | 7265 | 1,000 | -11,522 | 0,452 | -2,128 | 0,952 | 7061 | 1,000 |
| sample_00106 | -2,692 | 0,251 | 6166 | 1,000 | 9,103 | 0,461 | 7496 | 1,000 | -11,795 | 0,543 | -2,301 | 0,997 | 8325 | 1,000 |
| sample_00107 | -2,294 | 0,261 | 6784 | 1,000 | 8,980 | 0,423 | 10913 | 1,000 | -11,274 | 0,505 | -2,163 | 0,957 | 9361 | 1,000 |
| sample_00108 | -3,792 | 0,229 | 2290 | 1,000 | 9,068 | 0,526 | 5760 | 1,000 | -12,860 | 0,631 | -2,438 | 1,068 | 6508 | 1,000 |
| sample_00109 | -2,511 | 0,520 | 3720 | 1,000 | 8,378 | 0,439 | 4305 | 1,001 | -10,889 | 0,735 | -4,074 | 1,324 | 2748 | 1,000 |
| sample_00110 | -2,362 | 0,394 | 7416 | 1,000 | 8,493 | 0,529 | 7428 | 1,000 | -10,855 | 0,736 | -2,683 | 1,151 | 9270 | 1,000 |
| sample_00111 | -1,968 | 0,237 | 6345 | 1,000 | 9,064 | 0,352 | 9377 | 1,000 | -11,032 | 0,438 | -1,929 | 0,885 | 7894 | 1,000 |
| sample_00112 | -2,098 | 0,196 | 5695 | 1,000 | 9,191 | 0,294 | 7724 | 1,000 | -11,289 | 0,373 | -1,971 | 0,889 | 8753 | 1,000 |
| sample_00113 | -1,983 | 0,279 | 7702 | 1,000 | 8,948 | 0,383 | 10446 | 1,000 | -10,931 | 0,495 | -2,031 | 0,904 | 10991 | 1,000 |
| sample_00114 | -2,090 | 0,177 | 5739 | 1,000 | 9,228 | 0,352 | 9640 | 1,000 | -11,318 | 0,406 | -1,893 | 0,888 | 8851 | 1,000 |
| sample_00115 | -2,163 | 0,191 | 5003 | 1,000 | 9,324 | 0,386 | 7093 | 1,000 | -11,487 | 0,440 | -1,968 | 0,924 | 7116 | 1,001 |
| sample_00116 | -2,078 | 0,163 | 5714 | 1,000 | 9,268 | 0,355 | 8475 | 1,000 | -11,346 | 0,399 | -1,936 | 0,901 | 8589 | 1,000 |
| sample_00117 | -3,222 | 0,300 | 3613 | 1,002 | 9,283 | 0,468 | 7038 | 1,000 | -12,505 | 0,619 | -2,322 | 1,001 | 8618 | 1,000 |
| sample_00118 | -3,096 | 0,230 | 3977 | 1,000 | 9,359 | 0,427 | 7141 | 1,000 | -12,455 | 0,540 | -2,254 | 0,973 | 7220 | 1,000 |
| sample_00119 | -2,763 | 0,156 | 4991 | 1,000 | 9,475 | 0,359 | 5600 | 1,000 | -12,238 | 0,415 | -2,089 | 0,919 | 6594 | 1,000 |
| sample_00120 | -2,579 | 0,244 | 5104 | 1,000 | 9,035 | 0,406 | 8775 | 1,000 | -11,615 | 0,494 | -2,216 | 0,986 | 9368 | 1,001 |
| sample_00121 | -2,367 | 0,209 | 5185 | 1,000 | 9,367 | 0,340 | 8681 | 1,000 | -11,734 | 0,425 | -1,961 | 0,911 | 7323 | 1,000 |
| sample_00122 | -3,042 | 0,328 | 6956 | 1,000 | 9,217 | 0,493 | 10385 | 1,000 | -12,259 | 0,625 | -2,322 | 1,021 | 9399 | 1,000 |
| sample_00123 | -3,004 | 0,439 | 4124 | 1,001 | 9,213 | 0,501 | 7078 | 1,000 | -12,217 | 0,739 | -2,347 | 1,022 | 9253 | 1,000 |
| sample_00124 | -2,716 | 0,210 | 6081 | 1,000 | 9,292 | 0,400 | 9765 | 1,000 | -12,008 | 0,488 | -2,239 | 0,961 | 10454 | 1,000 |
| sample_00125 | -2,962 | 0,342 | 6050 | 1,000 | 9,205 | 0,494 | 10479 | 1,000 | -12,167 | 0,636 | -2,339 | 1,017 | 11857 | 1,000 |

**Table S24.** Ratios of the predicted trait values according to the linear models for cell size, gross speed and turning angles for evolved predators divided by ancestral predators. Prey and predator densities represent 2%, 50% and 95% quantiles of the observed prey and predator densities during the experiment. Note that for cell size, predictions are independent of prey species, and for movement speed, independent of prey densities.

| *Prey species* | *Prey evolution* | *Log (Prey density)* | *Log (Predator density)* | *size_ratio* | *speed_ratio* | *turning_ratio* |
|---|---|---|---|---|---|---|
| *E. coli* | Ancestor | 16,631 | 1,979 | 1,266 | 0,971 | 0,924 |
| *E. coli* | Ancestor | 16,631 | 6,066 | 1,235 | 1,042 | 0,965 |
| *E. coli* | Ancestor | 16,631 | 9,936 | 1,194 | 1,140 | 1,006 |
| *E. coli* | Ancestor | 19,021 | 1,979 | 1,110 | 0,971 | 0,924 |
| *E. coli* | Ancestor | 19,021 | 6,066 | 1,054 | 1,042 | 0,965 |
| *E. coli* | Ancestor | 19,021 | 9,936 | 0,994 | 1,140 | 1,006 |
| *E. coli* | Ancestor | 20,807 | 1,979 | 1,017 | 0,971 | 0,924 |
| *E. coli* | Ancestor | 20,807 | 6,066 | 0,963 | 1,042 | 0,965 |
| *E. coli* | Ancestor | 20,807 | 9,936 | 0,913 | 1,140 | 1,006 |
| *E. coli* | Evolved | 16,631 | 1,979 | 1,281 | 0,972 | 0,924 |
| *E. coli* | Evolved | 16,631 | 6,066 | 1,251 | 1,043 | 0,965 |
| *E. coli* | Evolved | 16,631 | 9,936 | 1,210 | 1,164 | 1,006 |
| *E. coli* | Evolved | 19,021 | 1,979 | 1,115 | 0,972 | 0,924 |
| *E. coli* | Evolved | 19,021 | 6,066 | 1,057 | 1,043 | 0,965 |
| *E. coli* | Evolved | 19,021 | 9,936 | 0,994 | 1,164 | 1,006 |
| *E. coli* | Evolved | 20,807 | 1,979 | 1,018 | 0,972 | 0,924 |
| *E. coli* | Evolved | 20,807 | 6,066 | 0,962 | 1,043 | 0,965 |
| *E. coli* | Evolved | 20,807 | 9,936 | 0,909 | 1,164 | 1,006 |
| *J. lividum* | Ancestor | 16,631 | 1,979 | 1,266 | 0,965 | 0,924 |
| *J. lividum* | Ancestor | 16,631 | 6,066 | 1,235 | 1,054 | 0,965 |
| *J. lividum* | Ancestor | 16,631 | 9,936 | 1,194 | 1,194 | 1,006 |
| *J. lividum* | Ancestor | 19,021 | 1,979 | 1,110 | 0,965 | 0,924 |
| *J. lividum* | Ancestor | 19,021 | 6,066 | 1,054 | 1,054 | 0,965 |
| *J. lividum* | Ancestor | 19,021 | 9,936 | 0,994 | 1,194 | 1,006 |
| *J. lividum* | Ancestor | 20,807 | 1,979 | 1,017 | 0,965 | 0,924 |
| *J. lividum* | Ancestor | 20,807 | 6,066 | 0,963 | 1,054 | 0,965 |
| *J. lividum* | Ancestor | 20,807 | 9,936 | 0,913 | 1,194 | 1,006 |
| *J. lividum* | Evolved | 16,631 | 1,979 | 1,281 | 0,966 | 0,924 |
| *J. lividum* | Evolved | 16,631 | 6,066 | 1,251 | 1,057 | 0,965 |
| *J. lividum* | Evolved | 16,631 | 9,936 | 1,210 | 1,244 | 1,006 |
| *J. lividum* | Evolved | 19,021 | 1,979 | 1,115 | 0,966 | 0,924 |
| *J. lividum* | Evolved | 19,021 | 6,066 | 1,057 | 1,057 | 0,965 |
| *J. lividum* | Evolved | 19,021 | 9,936 | 0,994 | 1,244 | 1,006 |

| | | | | | | |
|---|---|---|---|---|---|---|
| *J. lividum* | Evolved | 20,807 | 1,979 | 1,018 | 0,966 | 0,924 |
| *J. lividum* | Evolved | 20,807 | 6,066 | 0,962 | 1,057 | 0,965 |
| *J. lividum* | Evolved | 20,807 | 9,936 | 0,909 | 1,244 | 1,006 |
| *S. capsulata* | Ancestor | 16,631 | 1,979 | 1,266 | 0,971 | 0,924 |
| *S. capsulata* | Ancestor | 16,631 | 6,066 | 1,235 | 1,042 | 0,965 |
| *S. capsulata* | Ancestor | 16,631 | 9,936 | 1,194 | 1,143 | 1,006 |
| *S. capsulata* | Ancestor | 19,021 | 1,979 | 1,110 | 0,971 | 0,924 |
| *S. capsulata* | Ancestor | 19,021 | 6,066 | 1,054 | 1,042 | 0,965 |
| *S. capsulata* | Ancestor | 19,021 | 9,936 | 0,994 | 1,143 | 1,006 |
| *S. capsulata* | Ancestor | 20,807 | 1,979 | 1,017 | 0,971 | 0,924 |
| *S. capsulata* | Ancestor | 20,807 | 6,066 | 0,963 | 1,042 | 0,965 |
| *S. capsulata* | Ancestor | 20,807 | 9,936 | 0,913 | 1,143 | 1,006 |
| *S. capsulata* | Evolved | 16,631 | 1,979 | 1,281 | 0,972 | 0,924 |
| *S. capsulata* | Evolved | 16,631 | 6,066 | 1,251 | 1,044 | 0,965 |
| *S. capsulata* | Evolved | 16,631 | 9,936 | 1,210 | 1,168 | 1,006 |
| *S. capsulata* | Evolved | 19,021 | 1,979 | 1,115 | 0,972 | 0,924 |
| *S. capsulata* | Evolved | 19,021 | 6,066 | 1,057 | 1,044 | 0,965 |
| *S. capsulata* | Evolved | 19,021 | 9,936 | 0,994 | 1,168 | 1,006 |
| *S. capsulata* | Evolved | 20,807 | 1,979 | 1,018 | 0,972 | 0,924 |
| *S. capsulata* | Evolved | 20,807 | 6,066 | 0,962 | 1,044 | 0,965 |
| *S. capsulata* | Evolved | 20,807 | 9,936 | 0,909 | 1,168 | 1,006 |
| *B. diminuta* | Ancestor | 16,631 | 1,979 | 1,266 | 0,973 | 0,924 |
| *B. diminuta* | Ancestor | 16,631 | 6,066 | 1,235 | 1,039 | 0,965 |
| *B. diminuta* | Ancestor | 16,631 | 9,936 | 1,194 | 1,130 | 1,006 |
| *B. diminuta* | Ancestor | 19,021 | 1,979 | 1,110 | 0,973 | 0,924 |
| *B. diminuta* | Ancestor | 19,021 | 6,066 | 1,054 | 1,039 | 0,965 |
| *B. diminuta* | Ancestor | 19,021 | 9,936 | 0,994 | 1,130 | 1,006 |
| *B. diminuta* | Ancestor | 20,807 | 1,979 | 1,017 | 0,973 | 0,924 |
| *B. diminuta* | Ancestor | 20,807 | 6,066 | 0,963 | 1,039 | 0,965 |
| *B. diminuta* | Ancestor | 20,807 | 9,936 | 0,913 | 1,130 | 1,006 |
| *B. diminuta* | Evolved | 16,631 | 1,979 | 1,281 | 0,974 | 0,924 |
| *B. diminuta* | Evolved | 16,631 | 6,066 | 1,251 | 1,041 | 0,965 |
| *B. diminuta* | Evolved | 16,631 | 9,936 | 1,210 | 1,151 | 1,006 |
| *B. diminuta* | Evolved | 19,021 | 1,979 | 1,115 | 0,974 | 0,924 |
| *B. diminuta* | Evolved | 19,021 | 6,066 | 1,057 | 1,041 | 0,965 |
| *B. diminuta* | Evolved | 19,021 | 9,936 | 0,994 | 1,151 | 1,006 |
| *B. diminuta* | Evolved | 20,807 | 1,979 | 1,018 | 0,974 | 0,924 |
| *B. diminuta* | Evolved | 20,807 | 6,066 | 0,962 | 1,041 | 0,965 |
| *B. diminuta* | Evolved | 20,807 | 9,936 | 0,909 | 1,151 | 1,006 |

| | | | | | | |
|---|---|---|---|---|---|---|
| *P. fluorescens* | Ancestor | 16,631 | 1,979 | 1,266 | 0,970 | 0,924 |
| *P. fluorescens* | Ancestor | 16,631 | 6,066 | 1,235 | 1,044 | 0,965 |
| *P. fluorescens* | Ancestor | 16,631 | 9,936 | 1,194 | 1,150 | 1,006 |
| *P. fluorescens* | Ancestor | 19,021 | 1,979 | 1,110 | 0,970 | 0,924 |
| *P. fluorescens* | Ancestor | 19,021 | 6,066 | 1,054 | 1,044 | 0,965 |
| *P. fluorescens* | Ancestor | 19,021 | 9,936 | 0,994 | 1,150 | 1,006 |
| *P. fluorescens* | Ancestor | 20,807 | 1,979 | 1,017 | 0,970 | 0,924 |
| *P. fluorescens* | Ancestor | 20,807 | 6,066 | 0,963 | 1,044 | 0,965 |
| *P. fluorescens* | Ancestor | 20,807 | 9,936 | 0,913 | 1,150 | 1,006 |
| *P. fluorescens* | Evolved | 16,631 | 1,979 | 1,281 | 0,971 | 0,924 |
| *P. fluorescens* | Evolved | 16,631 | 6,066 | 1,251 | 1,046 | 0,965 |
| *P. fluorescens* | Evolved | 16,631 | 9,936 | 1,210 | 1,178 | 1,006 |
| *P. fluorescens* | Evolved | 19,021 | 1,979 | 1,115 | 0,971 | 0,924 |
| *P. fluorescens* | Evolved | 19,021 | 6,066 | 1,057 | 1,046 | 0,965 |
| *P. fluorescens* | Evolved | 19,021 | 9,936 | 0,994 | 1,178 | 1,006 |
| *P. fluorescens* | Evolved | 20,807 | 1,979 | 1,018 | 0,971 | 0,924 |
| *P. fluorescens* | Evolved | 20,807 | 6,066 | 0,962 | 1,046 | 0,965 |
| *P. fluorescens* | Evolved | 20,807 | 9,936 | 0,909 | 1,178 | 1,006 |
| *C. testosteroni* | Ancestor | 16,631 | 1,979 | 1,266 | 0,971 | 0,924 |
| *C. testosteroni* | Ancestor | 16,631 | 6,066 | 1,235 | 1,041 | 0,965 |
| *C. testosteroni* | Ancestor | 16,631 | 9,936 | 1,194 | 1,139 | 1,006 |
| *C. testosteroni* | Ancestor | 19,021 | 1,979 | 1,110 | 0,971 | 0,924 |
| *C. testosteroni* | Ancestor | 19,021 | 6,066 | 1,054 | 1,041 | 0,965 |
| *C. testosteroni* | Ancestor | 19,021 | 9,936 | 0,994 | 1,139 | 1,006 |
| *C. testosteroni* | Ancestor | 20,807 | 1,979 | 1,017 | 0,971 | 0,924 |
| *C. testosteroni* | Ancestor | 20,807 | 6,066 | 0,963 | 1,041 | 0,965 |
| *C. testosteroni* | Ancestor | 20,807 | 9,936 | 0,913 | 1,139 | 1,006 |
| *C. testosteroni* | Evolved | 16,631 | 1,979 | 1,281 | 0,973 | 0,924 |
| *C. testosteroni* | Evolved | 16,631 | 6,066 | 1,251 | 1,043 | 0,965 |
| *C. testosteroni* | Evolved | 16,631 | 9,936 | 1,210 | 1,162 | 1,006 |
| *C. testosteroni* | Evolved | 19,021 | 1,979 | 1,115 | 0,973 | 0,924 |
| *C. testosteroni* | Evolved | 19,021 | 6,066 | 1,057 | 1,043 | 0,965 |
| *C. testosteroni* | Evolved | 19,021 | 9,936 | 0,994 | 1,162 | 1,006 |
| *C. testosteroni* | Evolved | 20,807 | 1,979 | 1,018 | 0,973 | 0,924 |
| *C. testosteroni* | Evolved | 20,807 | 6,066 | 0,962 | 1,043 | 0,965 |
| *C. testosteroni* | Evolved | 20,807 | 9,936 | 0,909 | 1,162 | 1,006 |
| *S. marcescens* | Ancestor | 16,631 | 1,979 | 1,266 | 0,966 | 0,924 |
| *S. marcescens* | Ancestor | 16,631 | 6,066 | 1,235 | 1,051 | 0,965 |
| *S. marcescens* | Ancestor | 16,631 | 9,936 | 1,194 | 1,181 | 1,006 |

| | | | | | | |
|---|---|---|---|---|---|---|
| *S. marcescens* | Ancestor | 19,021 | 1,979 | 1,110 | 0,966 | 0,924 |
| *S. marcescens* | Ancestor | 19,021 | 6,066 | 1,054 | 1,051 | 0,965 |
| *S. marcescens* | Ancestor | 19,021 | 9,936 | 0,994 | 1,181 | 1,006 |
| *S. marcescens* | Ancestor | 20,807 | 1,979 | 1,017 | 0,966 | 0,924 |
| *S. marcescens* | Ancestor | 20,807 | 6,066 | 0,963 | 1,051 | 0,965 |
| *S. marcescens* | Ancestor | 20,807 | 9,936 | 0,913 | 1,181 | 1,006 |
| *S. marcescens* | Evolved | 16,631 | 1,979 | 1,281 | 0,968 | 0,924 |
| *S. marcescens* | Evolved | 16,631 | 6,066 | 1,251 | 1,054 | 0,965 |
| *S. marcescens* | Evolved | 16,631 | 9,936 | 1,210 | 1,223 | 1,006 |
| *S. marcescens* | Evolved | 19,021 | 1,979 | 1,115 | 0,968 | 0,924 |
| *S. marcescens* | Evolved | 19,021 | 6,066 | 1,057 | 1,054 | 0,965 |
| *S. marcescens* | Evolved | 19,021 | 9,936 | 0,994 | 1,223 | 1,006 |
| *S. marcescens* | Evolved | 20,807 | 1,979 | 1,018 | 0,968 | 0,924 |
| *S. marcescens* | Evolved | 20,807 | 6,066 | 0,962 | 1,054 | 0,965 |
| *S. marcescens* | Evolved | 20,807 | 9,936 | 0,909 | 1,223 | 1,006 |

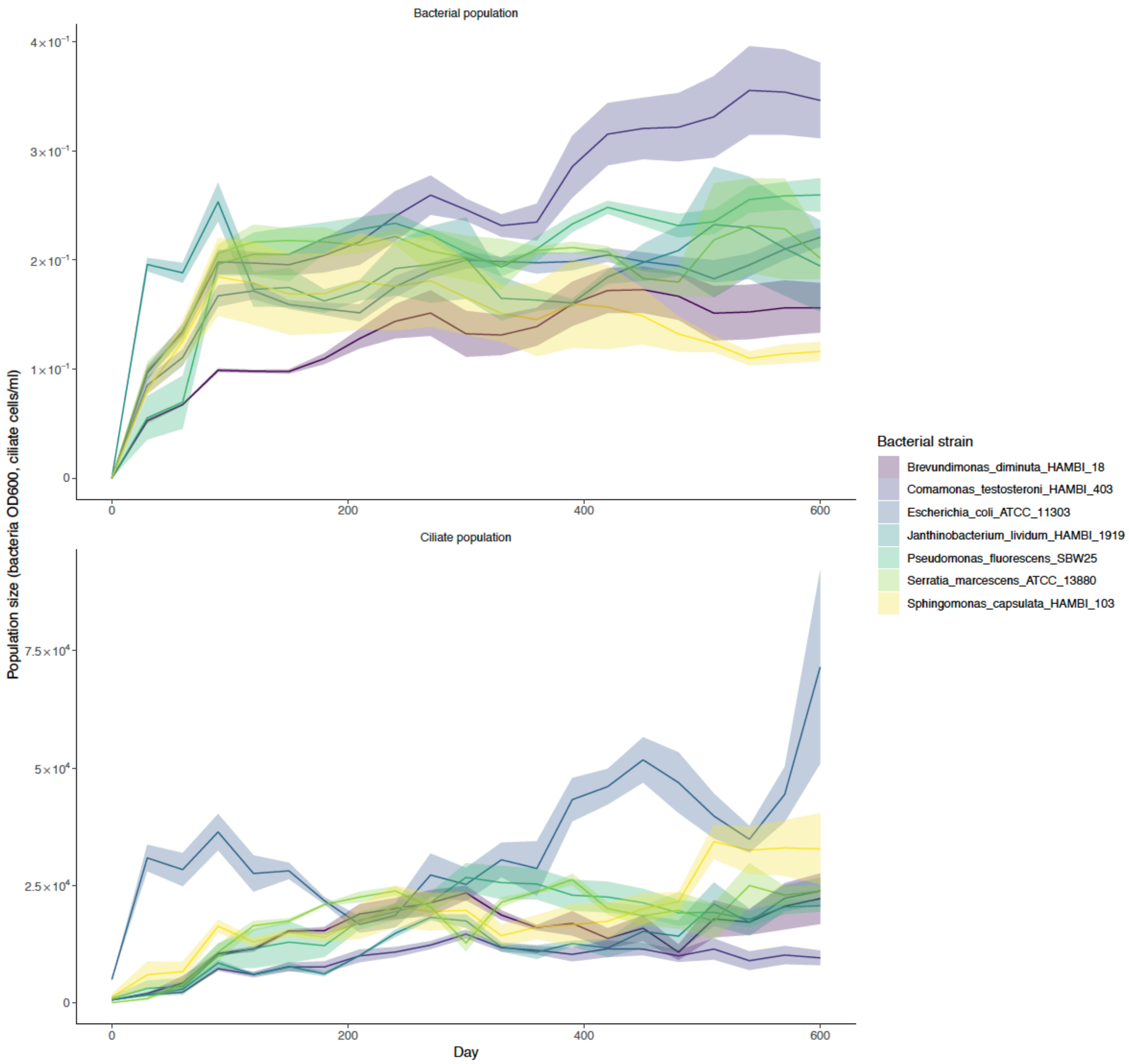


**Figure S1.** Bacterial and ciliate population size during first 20 months in long-term predator-prey coevolutionary experiment (mean ± s.e.m. smoothed over 90 day sliding window). The figure shows different population sizes for different bacterial species and an increasing trend over time.

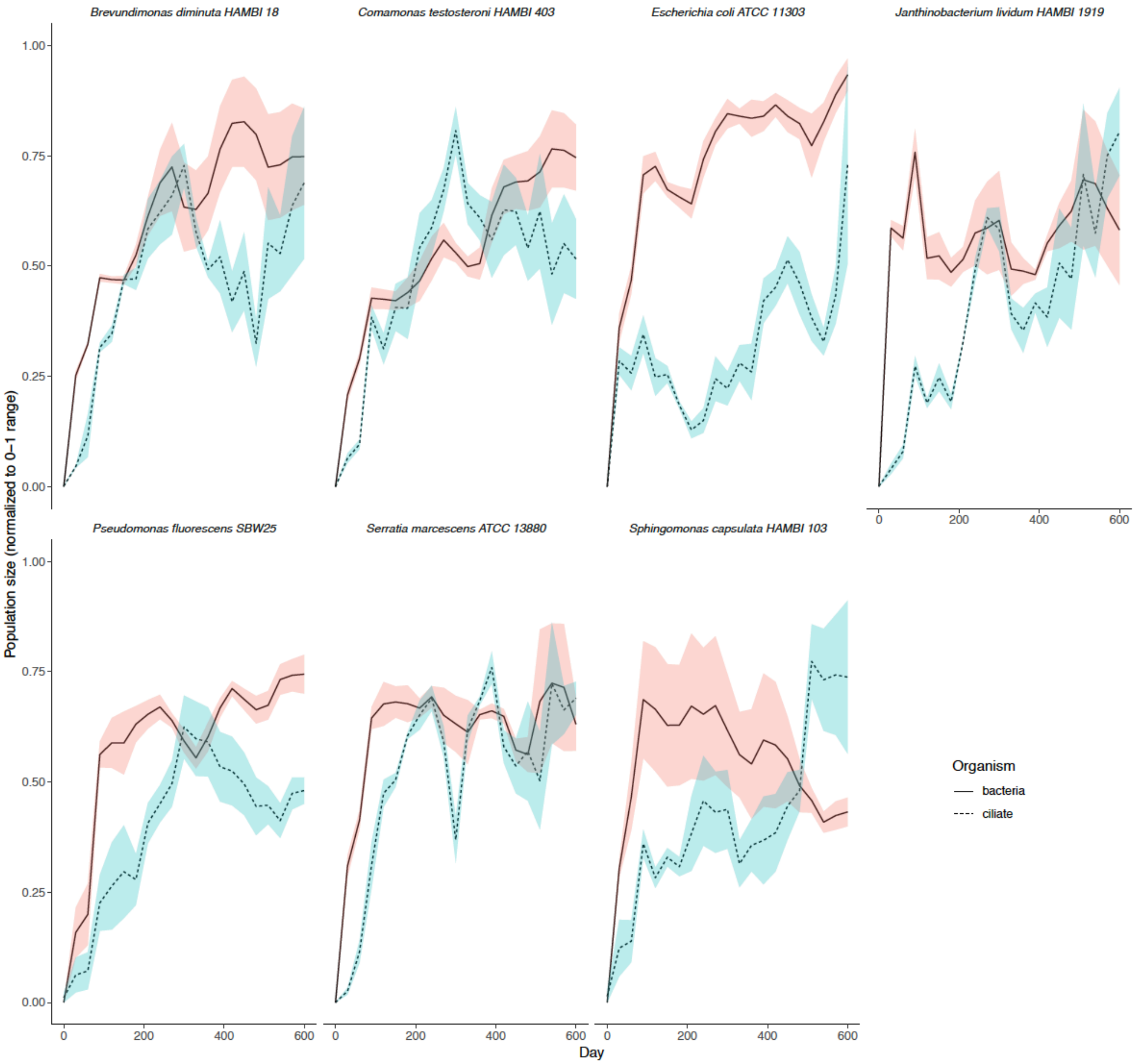


**Figure S2.** Bacteria-ciliate community dynamics during first 20 months in long-term predator-prey coevolutionary experiment (mean ± s.e.m. standardized to 0-1 range and smoothed over 90 day sliding window). The figure shows a negative association between bacterial and ciliate population size as well as differences in community dynamics depending on the prey (bacterial) species.

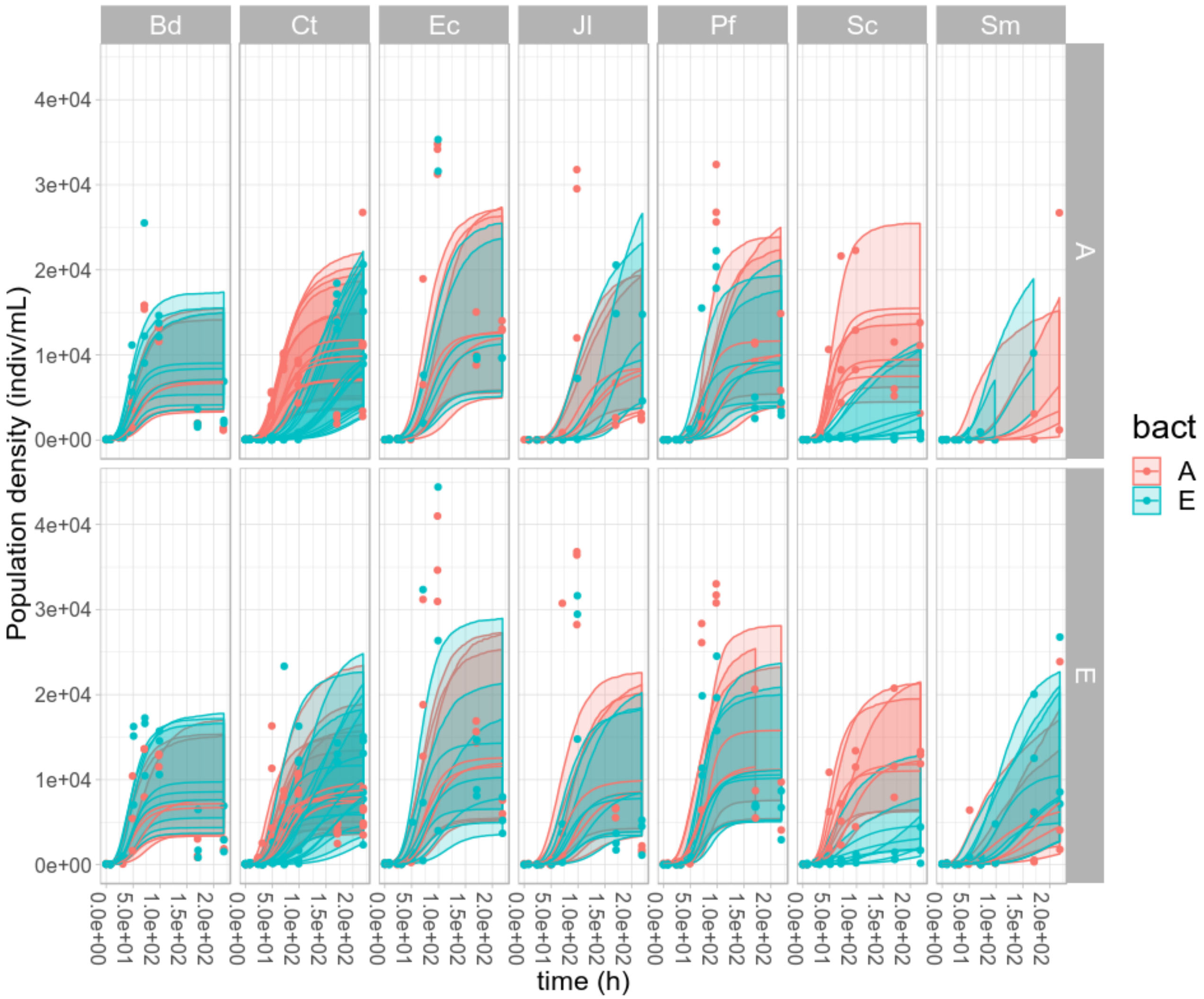


**Figure S3.** Beverton-Holt continuous-time growth models for ciliate population density (mean ± 95 % confidence intervals). A = ancestral ciliate; E = evolved ciliate; Bd = *Brevundimonas diminuta* HAMBI 18; Ct = *Comamonas testosteroni* HAMBI 403; Ec = *Escherichia coli* ATCC 11303; Jl = *Janthinobacterium lividum* HAMBI 1919; Pf = *Pseudomonas fluorescens* SBW25; Sc = *Sphingomonas capsulata* HAMBI 103; Sm = *Serratia marcescens* ATCC 13880.

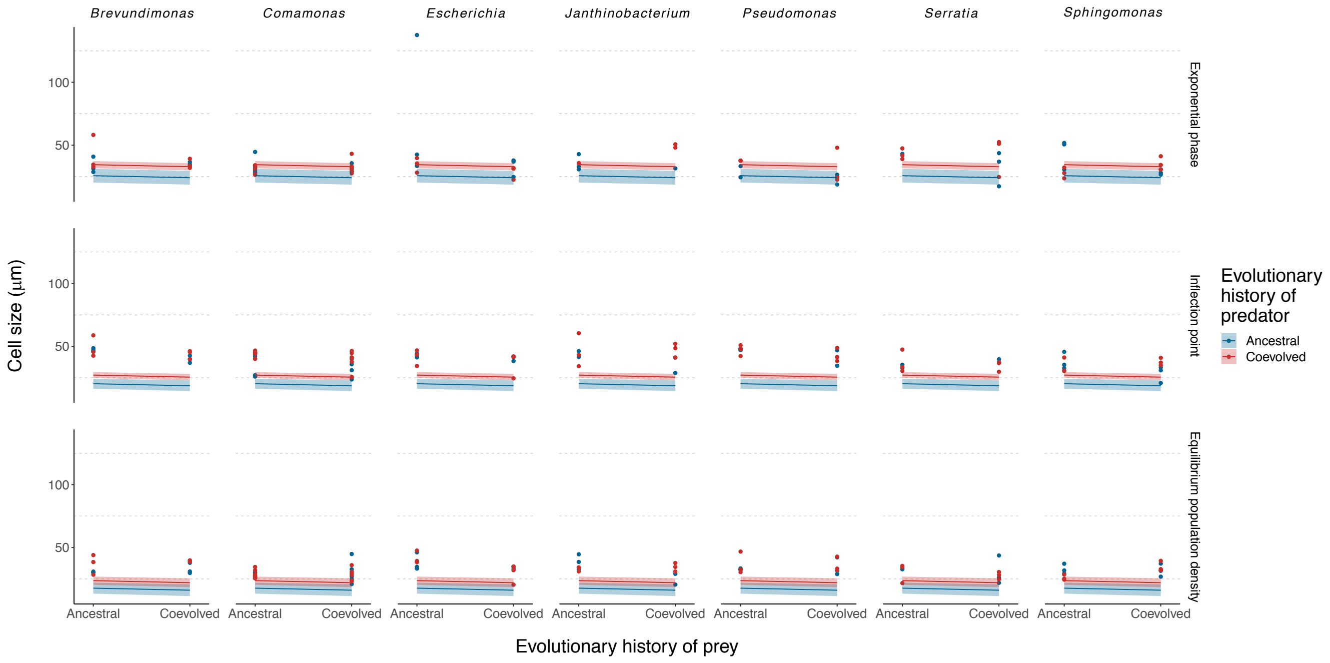


**Figure S4.** Reaction norms showing effect of predator-prey coevolution on cell size of predator at low (5 % quantile) prey density and three different predator densities (data points with linear model estimate ± 95 % confidence intervals.; N = 3 except 6 for *Comamonas*). Predator densities have been taken from different growth phases estimated using Beverton-Holt population models. The reaction norms for predators (one strain of the ciliate *Tetrahymena thermophila*) feeding on ancestral or coevolved prey (seven bacterial strains indicated by genus name) are depicted separately for ancestral and coevolved predators (color coding). Predators coevolved with a particular prey taxon have always been coupled with ancestral or coevolved populations of the same taxon, while the ancestral predator is the same for all prey taxa.

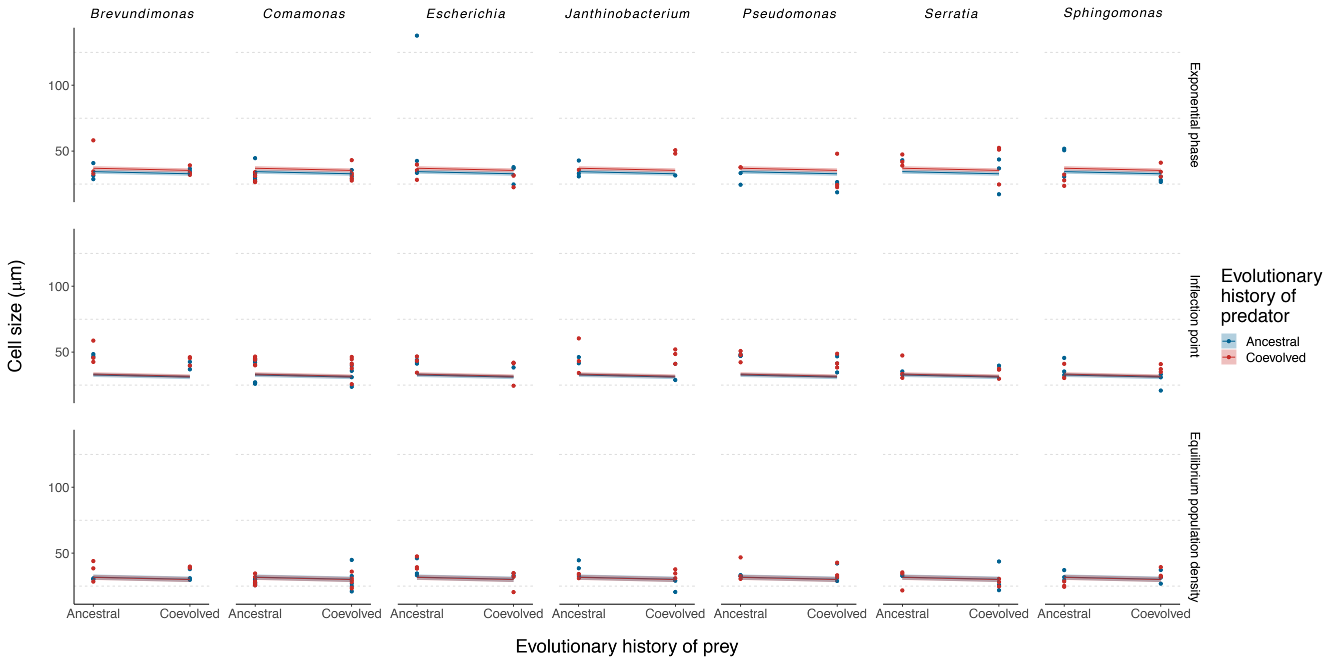


**Figure S5.** Reaction norms showing effect of predator-prey coevolution on cell size of predator at medium (50 % quantile) prey density and three different predator densities (data points with linear model estimate ± 95 % confidence intervals.; N = 3 except 6 for *Comamonas*). Predator densities have been taken from different growth phases estimated using Beverton-Holt population models. The reaction norms for predators (one strain of the ciliate *Tetrahymena thermophila*) feeding on ancestral or coevolved prey (seven bacterial strains indicated by genus name) are depicted separately for ancestral and coevolved predators (color coding). Predators coevolved with a particular prey taxon have always been coupled with ancestral or coevolved populations of the same taxon, while the ancestral predator is the same for all prey taxa.

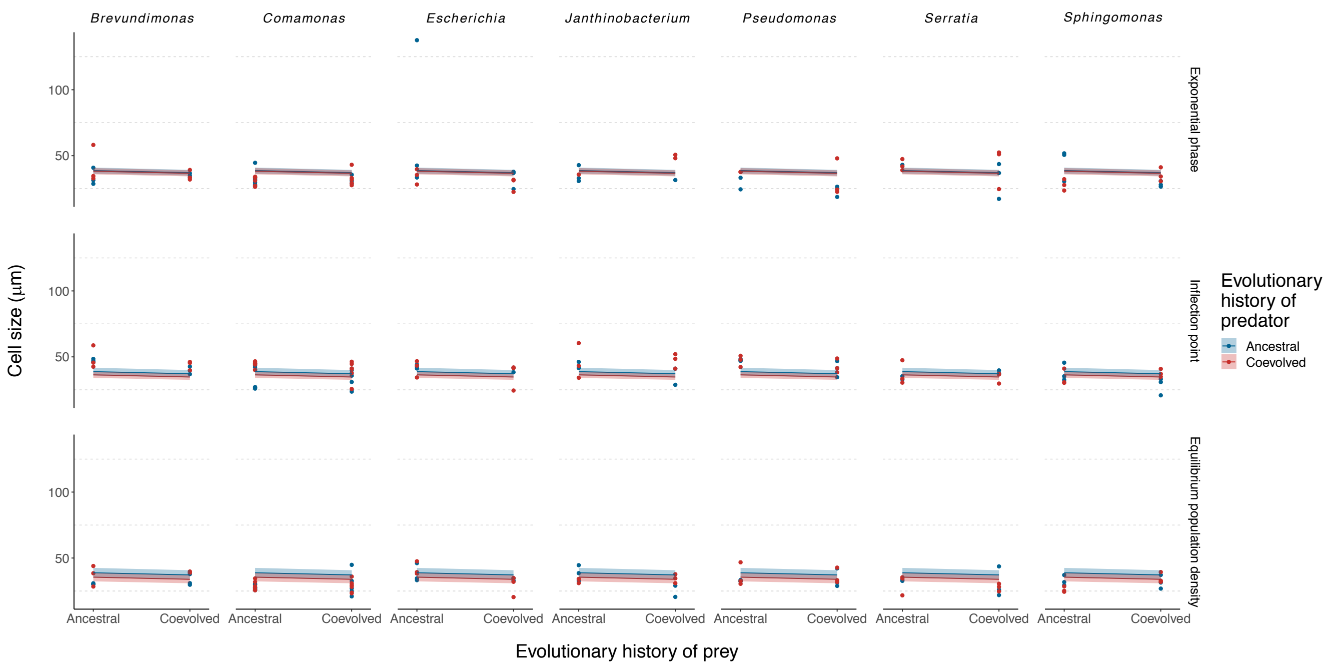


**Figure S6.** Reaction norms showing effect of predator-prey coevolution on cell size of predator at high (95 % quantile) prey density and three different predator densities (data points with linear model estimate ± 95 % confidence intervals.; N = 3 except 6 for *Comamonas*). Predator densities have been taken from different growth phases estimated using Beverton-Holt population models. The reaction norms for predators (one strain of the ciliate *Tetrahymena thermophila*) feeding on ancestral or coevolved prey (seven bacterial strains indicated by genus name) are depicted separately for ancestral and coevolved predators (color coding). Predators coevolved with a particular prey taxon have always been coupled with ancestral or coevolved populations of the same taxon, while the ancestral predator is the same for all prey taxa.

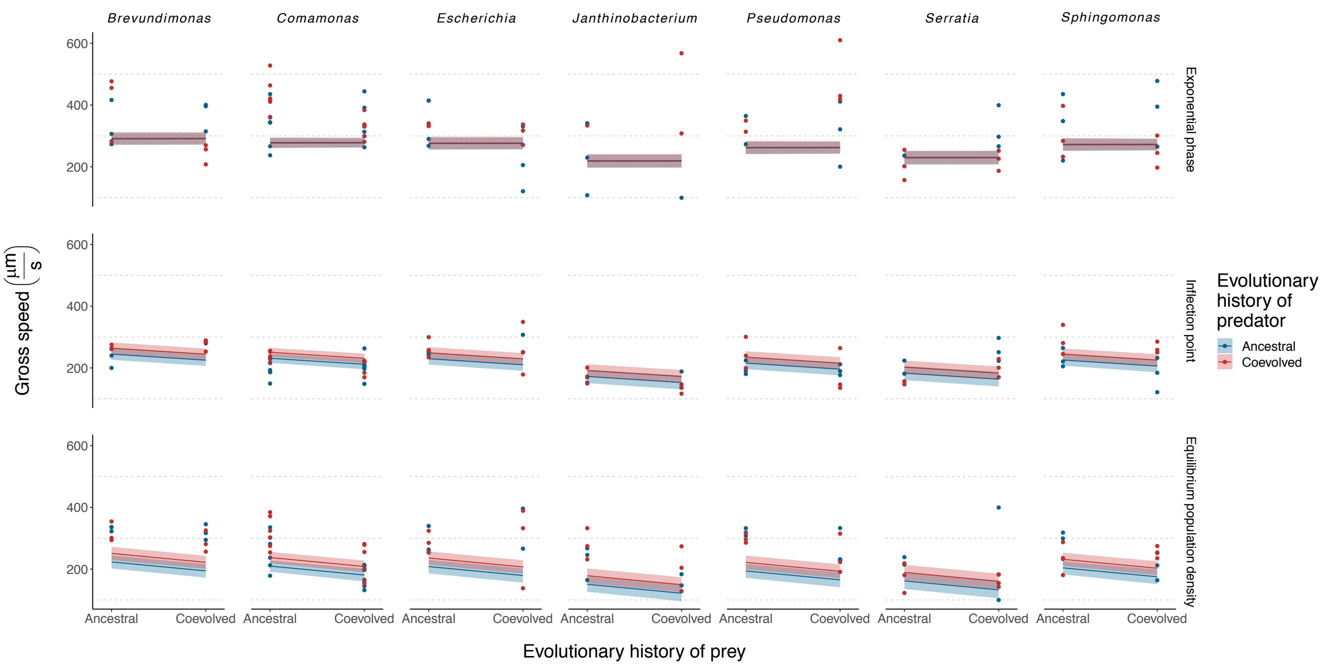


**Figure S7.** Reaction norms showing effect of predator-prey coevolution on gross speed of predator at medium (50 % quantile) prey density and three different predator densities (data points with linear model estimate ± 95 % confidence intervals.; N = 3 except 6 for *Comamonas*). Since the statistical analysis did not show an effect of prey density on gross speed of the predator, only one prey density is plotted for speed unlike for cell size and turning angle distribution. Predator densities have been taken from different growth phases estimated using Beverton-Holt population models. The reaction norms for predators (one strain of the ciliate *Tetrahymena thermophila*) feeding on ancestral or coevolved prey (seven bacterial strains indicated by genus name) are depicted separately for ancestral and coevolved predators (color coding). Predators coevolved with a particular prey taxon have always been coupled with ancestral or coevolved populations of the same taxon, while the ancestral predator is the same for all prey taxa.

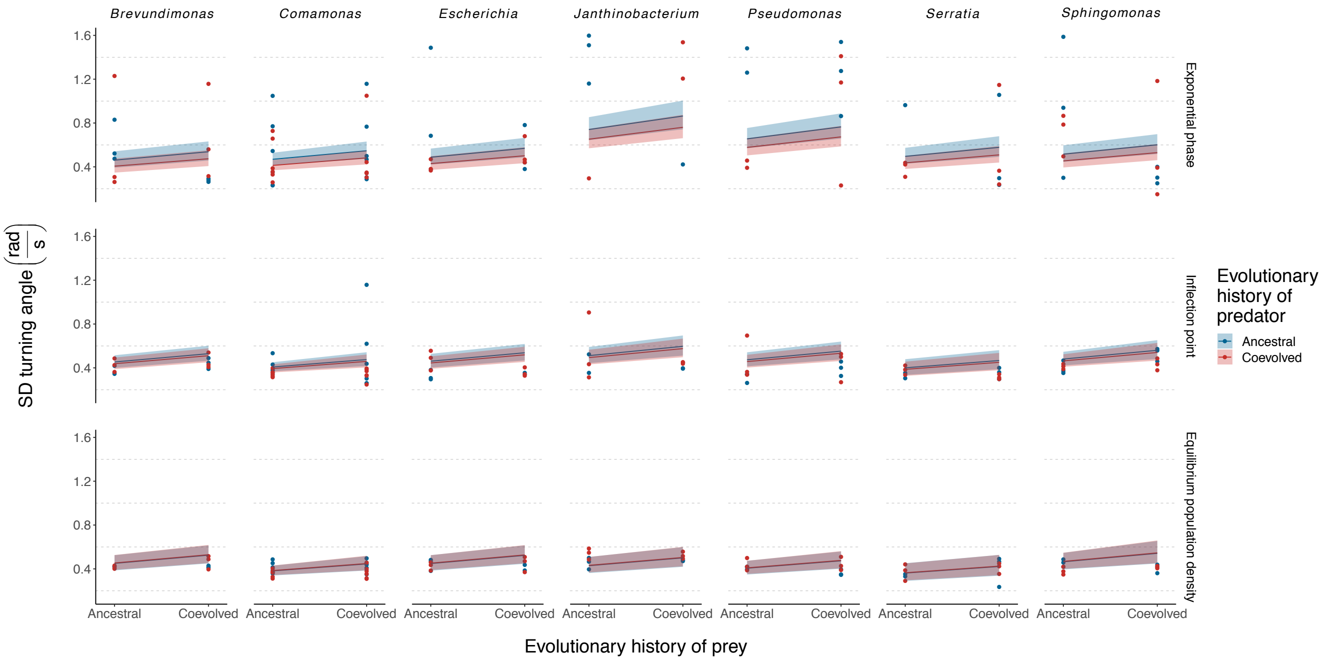


**Figure S8.** Reaction norms showing effect of predator-prey coevolution on turning angle distribution of predator at low (5 % quantile) prey density and three different predator densities (data points with linear model estimate ± 95 % confidence intervals.; N = 3 except 6 for *Comamonas*). Cell turning angle distribution (standard deviation, SD) is used as a proxy for directionality of cell movement which is higher at lower values. Predator densities have been taken from different growth phases estimated using Beverton-Holt population models. The reaction norms for predators (one strain of the ciliate *Tetrahymena thermophila*) feeding on ancestral or coevolved prey (seven bacterial strains indicated by genus name) are depicted separately for ancestral and coevolved predators (color coding). Predators coevolved with a particular prey taxon have always been coupled with ancestral or coevolved populations of the same taxon, while the ancestral predator is the same for all prey taxa.

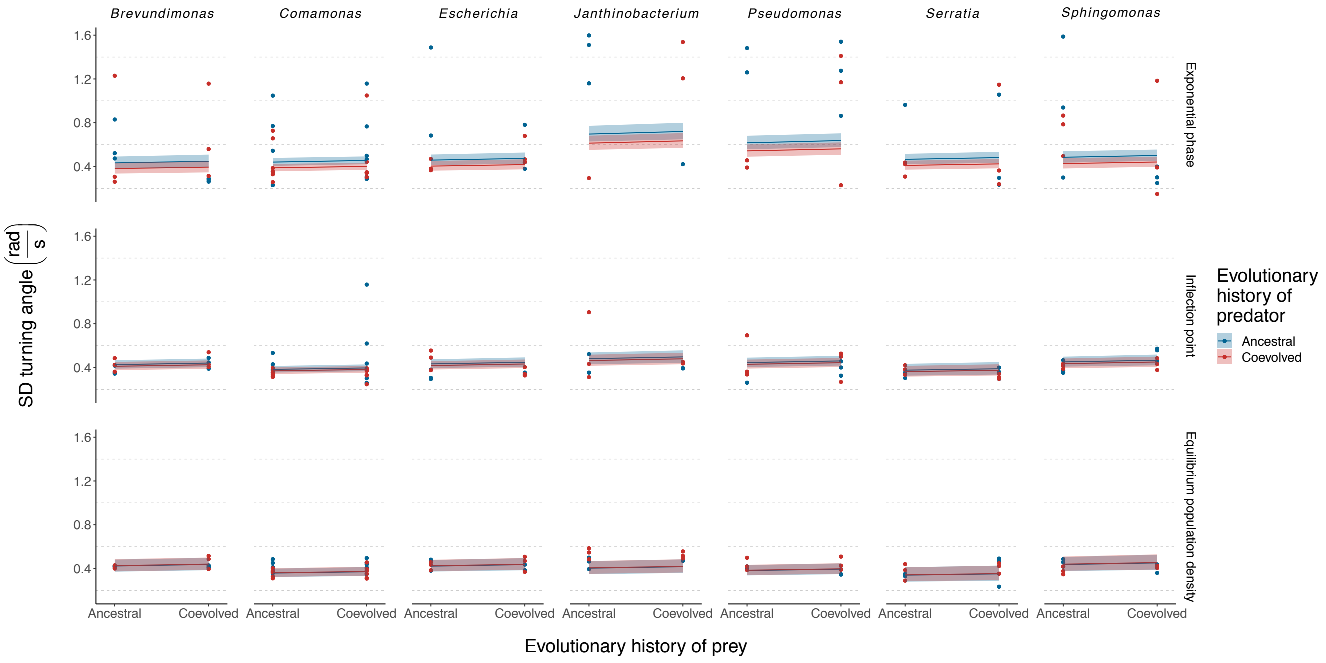


**Figure S9.** Reaction norms showing effect of predator-prey coevolution on turning angle distribution of predator at medium (50 % quantile) prey density and three different predator densities (data points with linear model estimate ± 95 % confidence intervals.; N = 3 except 6 for *Comamonas*). Cell turning angle distribution (standard deviation, SD) is used as a proxy for directionality of cell movement which is higher at lower values. Predator densities have been taken from different growth phases estimated using Beverton-Holt population models. The reaction norms for predators (one strain of the ciliate *Tetrahymena thermophila*) feeding on ancestral or coevolved prey (seven bacterial strains indicated by genus name) are depicted separately for ancestral and coevolved predators (color coding). Predators coevolved with a particular prey taxon have always been coupled with ancestral or coevolved populations of the same taxon, while the ancestral predator is the same for all prey taxa.

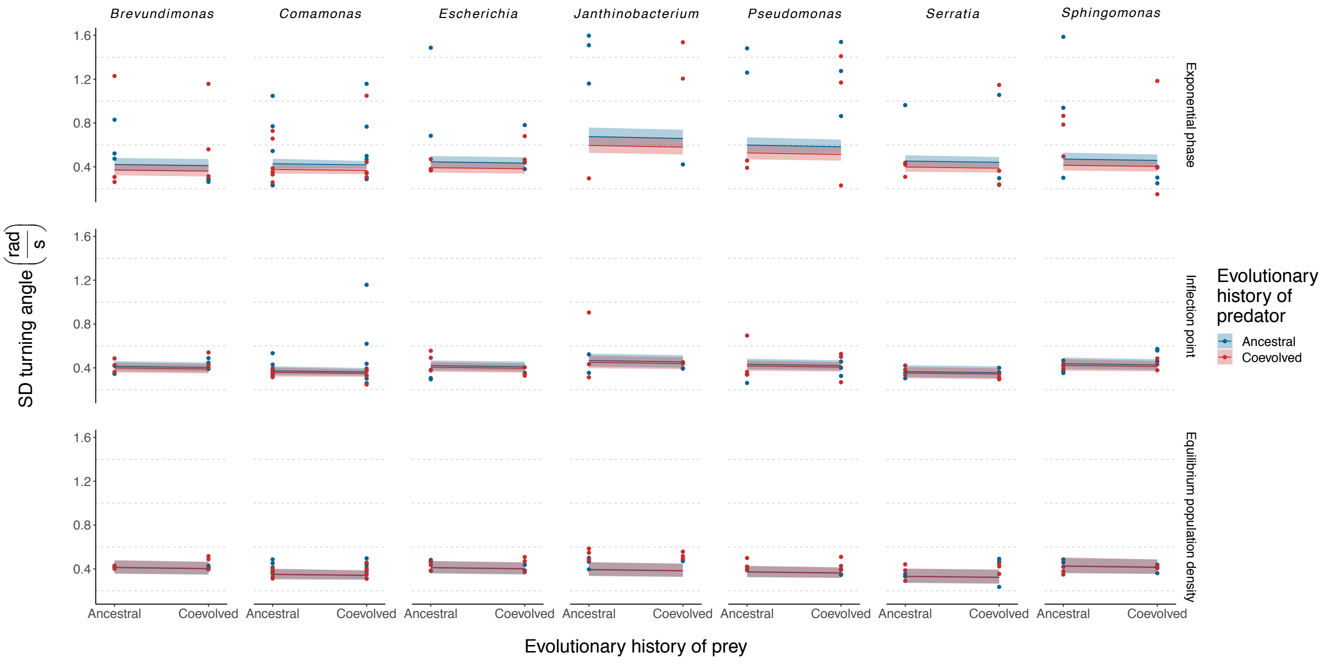


**Figure S10.** Reaction norms showing effect of predator-prey coevolution on turning angle distribution of predator at high (95 % quantile) prey density and three different predator densities (data points with linear model estimate ± 95 % confidence intervals.; N = 3 except 6 for *Comamonas*). Cell turning angle distribution (standard deviation, SD) is used as a proxy for directionality of cell movement which is higher at lower values. Predator densities have been taken from different growth phases estimated using Beverton-Holt population models. The reaction norms for predators (one strain of the ciliate *Tetrahymena thermophila*) feeding on ancestral or coevolved prey (seven bacterial strains indicated by genus name) are depicted separately for ancestral and coevolved predators (color coding). Predators coevolved with a particular prey taxon have always been coupled with ancestral or coevolved populations of the same taxon, while the ancestral predator is the same for all prey taxa.

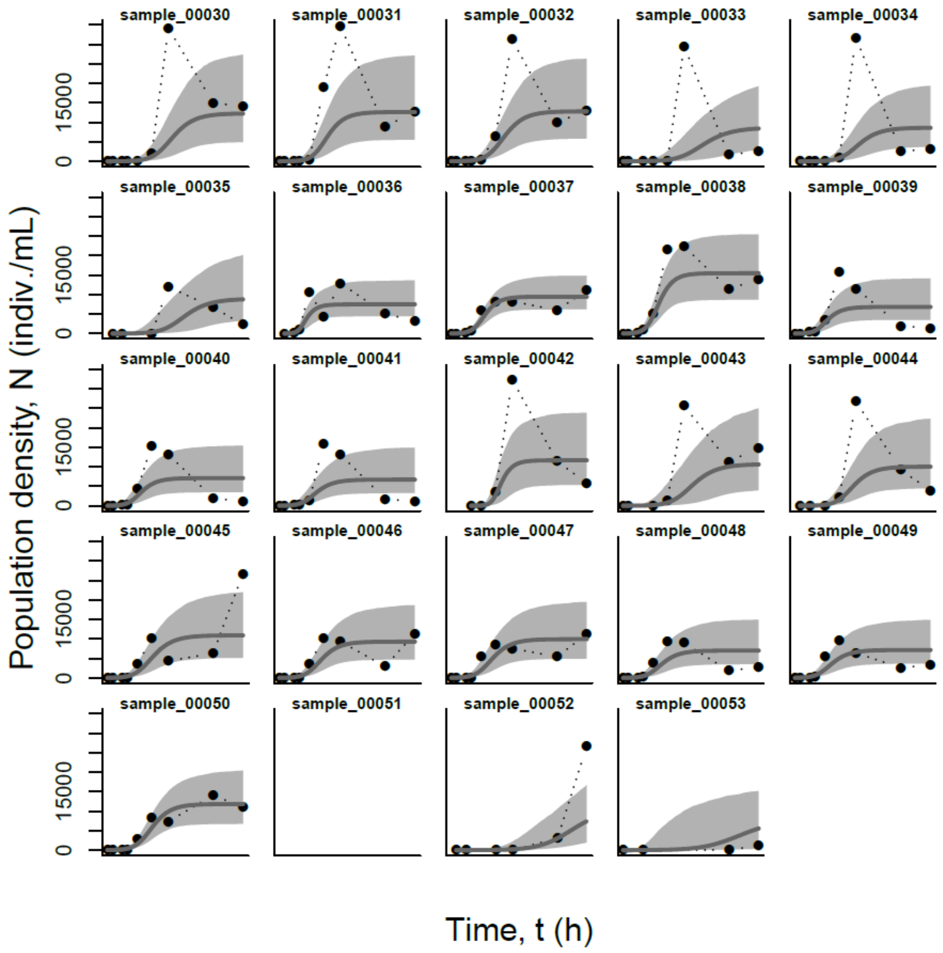


**Figure S11.** Beverton-Holt predictions for the individual populations (populations 1-24 (samples 30-53)). Dots represent population density measured using video analysis. The smooth grey line shows the mean model prediction, and shaded are the 95% probability interval of the model prediction.

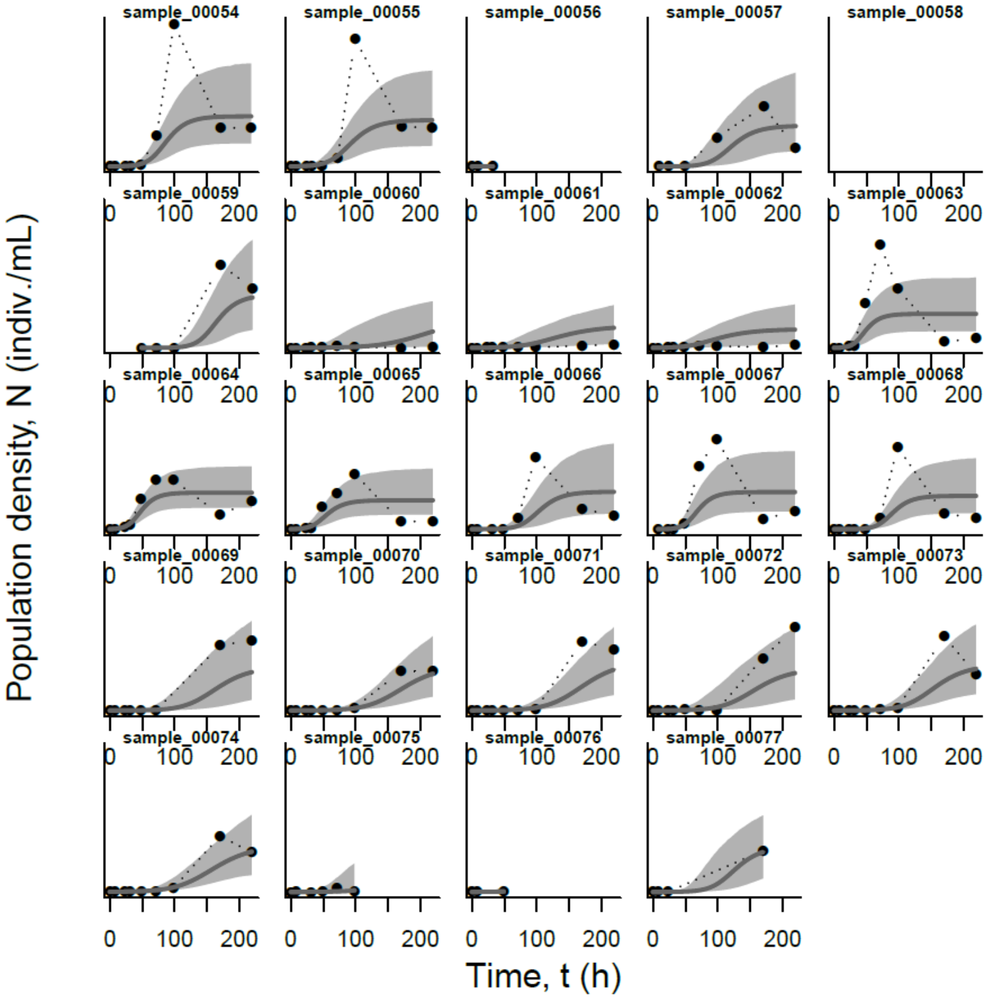


**Figure S12.** Beverton-Holt predictions for the individual populations (populations 25-48 (samples 54-77)). Dots represent population density measured using video analysis. The smooth grey line shows the mean model prediction, and shaded are the 95% probability interval of the model prediction.

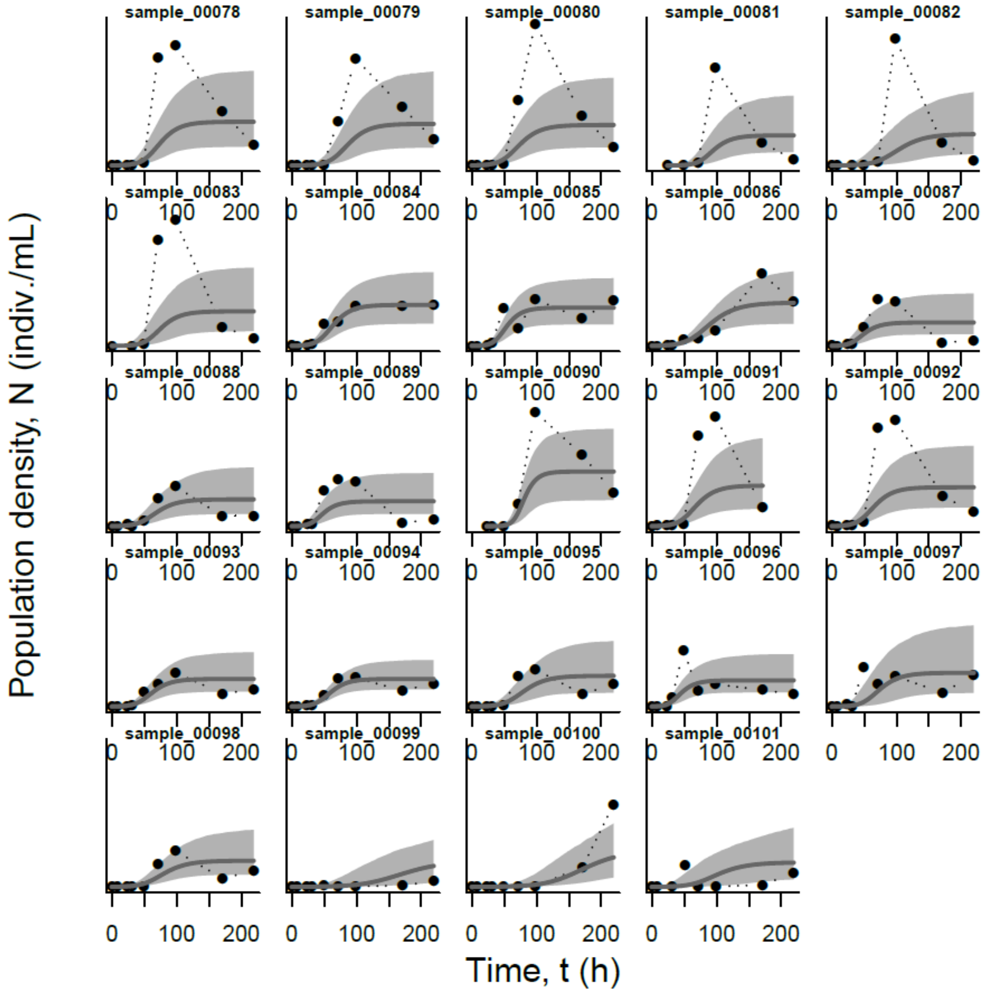


**Figure S13.** Beverton-Holt predictions for the individual populations (populations 49-72 (samples 78-101)). Dots represent population density measured using video analysis. The smooth grey line shows the mean model prediction, and shaded are the 95% probability interval of the model prediction.

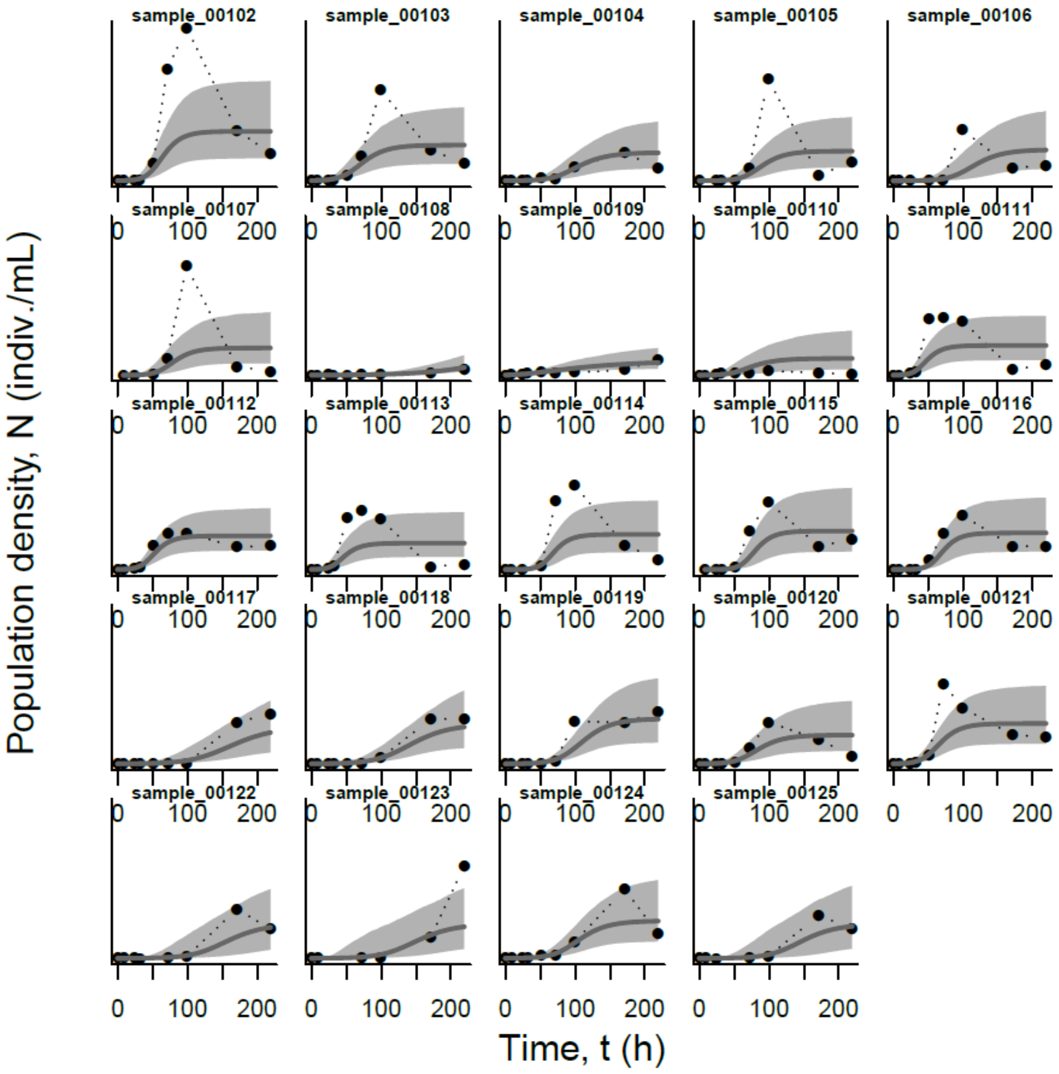


**Figure S14.** Beverton-Holt predictions for the individual populations (populations 49-72 (samples 78-101)). Dots represent population density measured using video analysis. The smooth grey line shows the mean model prediction, and shaded are the 95% probability interval of the model prediction.
